# Small-molecule modulators of TRMT2A decrease PolyQ aggregation and PolyQ-induced cell death

**DOI:** 10.1101/2021.05.03.442419

**Authors:** Michael A Margreiter, Monika Witzenberger, Yasmine Wasser, Elena Davydova, Robert Janowski, Carina Sobisch, Jonas Metz, Benedetta Poma, Oscar Palomino-Hernandez, Pardes Habib, Mirko Wagner, Thomas Carell, N Jon Shah, Jörg B Schulz, Dierk Niessing, Aaron Voigt, Giulia Rossetti

## Abstract

Polyglutamine (polyQ) diseases are characterized by an expansion of cytosine-adenine-guanine (CAG) trinucleotide repeats encoding for an uninterrupted prolonged polyQ tract. We previously identified TRMT2A as a strong modifier of polyQ-induced toxicity in an unbiased large-scale screen in *Drosophila melanogaster*. This work aimed at identifying and validating pharmacological TRMT2A inhibitors as treatment opportunities for polyQ diseases in humans. Computer-aided drug discovery was implemented to identify human TRMT2A inhibitors. Additionally, the crystal structure of one protein domain, the RNA recognition motif (RRM), was determined, and Biacore experiments with the RRM were performed. The identified molecules were validated for their potency to reduce polyQ aggregation and polyQ-induced cell death in human HEK293T cells and patient derived fibroblasts. Our work provides a first step towards pharmacological inhibition of this enzyme and indicates TRMT2A as a viable drug target for polyQ diseases.

## 1. Introduction

Polyglutamine (PolyQ) diseases are autosomal dominant inherited genetic disorders, except SMAX1/SBMA, which is recessive and linked to a mutation in the androgen receptor gene located on the X chromosome. PolyQ diseases are characterized by progressive neurodegeneration, leading to behavioral and physical impairments [1]. This disease family includes nine members: Spinobulbar muscular atrophy (SBMA or Kennedy’s disease) [2], Dentatorubral-pallidoluysian atrophy (Haw River syndrome) [3], Spinocerebellar ataxias 1, 2, 3 (Machado-Joseph disease), 6, 7, 17 [4] and Huntington’s disease (HD) [5]. Epidemiological data on HD, the most prevalent polyQ disease, reveals an incidence of 5-6 in 100.000 in the best-ascertained populations of Western Europe and North America [6]. All polyQ diseases feature an abnormally elongated CAG repeat expansion in the protein-coding region of the disease-linked gene [7]. As CAG encodes glutamine, the translation of mutant alleles results in a protein with an elongated, undisrupted polyQ stretch, eponymous for these diseases. The polyQ region itself is considered to be causative for these diseases since they usually form cytoplasmic and/or nuclear aggregates in affected neurons. Whether these aggregates contribute to neurodegeneration remains elusive; however, the appearance of polyQ aggregates coincides with neuronal dysfunction/loss [7]. The age of onset is inversely correlated with CAG tract length, while gene-specific differences between the various polyQ diseases have been reported [6]. Administration of antisense oligonucleotides (ASOs) to disease model animals suppressed expression of the disease-causing protein [8, 9]. Clinical trials with ASOs designed to degrade wild-type and mutant *Huntingtin* mRNA were started three years ago [10] but they were recently halted based on the drug’s potential benefit/risk profile for study participants [11]. Thus, the identification and targeting of novel CAG-disease modifiers remain a feasible therapeutic strategy.

In a high-throughput RNAi knock-down screen, tRNA methyltransferase 2 homolog A (TRMT2A) was identified as a novel modifier of polyQ-induced toxicity [12]. This protein predominantly localizes to the nucleus and converts uridine to 5-methyl uridine (m^5^U) at position 54 in tRNAs [13, 14]. The latter methylation is conserved and found from yeast to man, suggesting a pivotal role for this modification [15]. In *Escherichia coli*, the enzyme TrmA catalyzes the formation of m^5^U_54_ [16], while in *Saccharomyces cerevisiae*, the enzyme responsible for this modification is Trm2p [16]. TRMT2A is considered to represent the ortholog of *Saccharomyces cerevisiae* Trm2p [17] in mammals. Indeed, similar to Trm2p, TRMT2A was predicted to harbor an RNA recognition motif (RRM) at the N-terminus and a methyltransferase catalytic domain (CD) at the C-terminus [13]. In vertebrates, there is a second ortholog of Trm2p, TRMT2B. TRMT2A and TRMT2B are paralogs and exert similar functions. Whereas TRMT2A is responsible for the m^5^U_54_ methylation in the nucleus and cytoplasm, TRMT2B is specific for mitochondrial tRNAs [18]. Previous studies on *Escherichia coli* have shown that the presence of this methylation ensures the fidelity and efficiency of protein synthesis by stabilizing the three-dimensional structure of tRNA *in vitro* [19, 20]. So far, however, the biological function of the enzyme responsible for U_54_ methylation in mammalian species remains largely unexplored [13].

In this study, inhibitors of human TRMT2A were identified by *in silico* predictions. Two strategies of TRMT2A inhibition were followed, targeting the RRMs or the CD. In a second step, potential inhibitors were validated in human ‘HEK293T’ cells expressing polyQ peptides of pathogenic length for their ability to reduce cell-death and aggregation. Additionally, we used polyQ disease-patient-derived fibroblast for the most efficient inhibitors.

## 2. Material and Methods

### 2.1 Crystal Structure

#### 2.1.1 Cloning, expression and purification

Plasmid containing human the TRMT2A sequence was provided by Aaron Voigt. Human TRMT2A RRM (aa 69 – 147) was cloned with a cleavable SUMO-tag at the N-terminus and expressed in Rosetta *E. coli* cells in LB medium. After isopropyl-thiogalactoside (IPTG, 0.5mM) induction overnight at 18 °C, cells were harvested (4500 x g, 15 min, 4 °C) and flash-frozen with liquid nitrogen. All subsequent steps were carried out at 4 °C. Typically, 6 liters of culture were resuspended in lysis buffer (50 mM HEPES/NaOH pH 8.5, 500 mM NaCl, 20 mM imidazole, 0.5% (v/v) Tween, 2% (v/v) glycerol) supplemented with one tablet of EDTA-free protease inhibitor cocktail to a total volume of 50 ml. After sonication (4 x 6 min, amplitude 40%, output: 6), the lysate was clarified by centrifugation (40000 x g, 30 min), loaded onto 5 ml His-FFTrap column (GE Healthcare), equilibrated in His-A buffer (50 mM HEPES/NaOH pH 8.5, 500 mM NaCl, 20 mM imidazole), and washed with 10 column volumes (CV) of His-A buffer, followed by a 10 CV wash with His-B buffer (50 mM HEPES/NaOH pH 8.5, 2000 mM NaCl, 20 mM imidazole). SUMO tagged protein was eluted with a 10 CV gradient of His-A buffer and His-C elution buffer (50 mM HEPES/NaOH pH 8.5, 500 mM NaCl, 500 mM imidazole). For tag cleavage, the eluate was supplemented with 100 μg of HRV 3C protease (PreScission) and dialyzed against a dialysis buffer (50 mM HEPES/NaOH pH 7.5, 500 mM NaCl, 1 mM DTT) O/N. Cleaved protein was applied onto a His-Trap FF column, and the flow-through that contains cleaved protein was collected. This flow-through was concentrated and loaded onto a size exclusion chromatography column (Superdex 75 10/300 gl) equilibrated in SEC buffer (50 mM HEPES/NaOH pH 7.5, 500 mM NaCl). The purified protein was concentrated to 8 mg/ml by ultrafiltration. Protein concentrations were determined by measurement of the A_280_. Aliquots were flash-frozen in liquid nitrogen and stored at −80 °C.

#### 2.1.2 Crystallization, diffraction data collection, and processing

The crystallization experiments for the RRM domain were performed at the X-ray Crystallography Platform at Helmholtz Zentrum, München. Crystallization screening was done at 292 K using 8.0 mg/ml of protein with a nanodrop dispenser in sitting-drop 96-well plates and commercial screens. Crystals appeared after one week and were big enough for X-ray diffraction experiments. The initial data set for solving the structure was collected for a crystal grown in 0.22 M lithium sulfate, 0.1 M sodium acetate (pH 4.5), and 26% (w/v) PEG 6000 (Hampton Research PEG screen). However, the best data set used for refinement was collected for a crystal grown in 0.2 M cesium sulfate and 2.2 M ammonium sulfate (Hampton Research AmSO4 screen). For the X-ray diffraction experiments, the crystals were mounted in a nylon fiber loop and flash-cooled to 100 K in liquid nitrogen. Prior to freezing, the crystals were protected with 25% (v/v) ethylene glycol. Diffraction data were collected at 100 K on the P11 beamline (DESY, Hamburg) and PXIII beamline (SLS, Villigen). The diffraction data were indexed and integrated using *XDS [21]* and scaled using *SCALA [22, 23]*. Intensities were converted to structure-factor amplitudes using the program *TRUNCATE* [24]. Table S1 summarizes data collection and processing statistics.

#### 2.1.3 Structure determination and refinement

The structure of the RRM with a resolution of 2.02 Å was solved with the Auto-Rickshaw pipeline [25, 26] using the sequence as input. For the molecular replacement step followed by several cycles of automated model building and refinement, the Auto-Rickshaw pipeline invoked the following X-ray crystallography software: MORDA (Vagin and Lebedev; http://www.biomexsolutions.co.uk/morda/), CCP4 [27], SHELXE [28], BUCCANEER [29, 30], RESOLVE [31], REFMAC5 [32] and PHENIX [33]. Model rebuilding was performed in COOT [34]. Initial refinement was done with the 2.02 Å dataset in REFMAC5 [32] using the maximum-likelihood target function. Further refinement was done with another dataset with a higher resolution of 1.23 Å. The stereochemical analysis of the final model was done in PROCHECK [35] and MolProbity[35]. The final model is characterized by R/R_free_ factors of 12.60 / 18.90%. Atomic coordinates and structure factors for the 2.0 Å and 1.23 Å structures have been deposited to the Protein Data Bank under accession codes 7NTN and 7NTO, respectively.

### 2.2 In-silico Analyses of the Crystal Structure

#### 2.2.1 Binding Site Detection

Agglomerative Euclidean distance clustering on the crystallographic water (oxygen atom) clusters was done with Wolfram Mathematica 11.3. The X-ray structure was prepared with Maestro Schrödinger (Schrödinger Release 2016-2: Maestro, Schrödinger, LLC, New York, NY, 2019.) using default settings unless otherwise noted. Alignments were performed with ClustalW [36], while conservation analyses were performed with AL2CO [37]. For binding site detection, we used DoGSiteScorer [38], the transient site discovery, the machine learning-based CryptoSite [39] approach, and TRAPP [40]. TRAPP is a framework to generate and analyze large-scale secondary structure rearrangements on short molecular dynamics timescales, e.g., by disrupting sidechain atoms with short pulses (L-RIP). The system temperature is controlled by a Langevin thermostat to ensure a constant average temperature. In parallel, we investigated ligand-mediated conformational changes. We used an apolar/aromatic amino acid as a model for hydrophobic/aromatic ligand features and brought them into contact with the protein surface (RIPlig)[41]. Finally, we investigated putative allosteric effects in TRMT2A RRM with AlloPred [42].

#### 2.2.2. MD simulations

The protein was prepared using Maestro (Schrödinger Release 2016-2) and was visually inspected for correct protonation states. The partially occupied R137 with the side chain closer to the protein center was retained, as well as the crystallographic waters. To prepare the protein structure for the explicit solvent molecular dynamics simulations, tleap (AmberTools 17) was used, and the AMBER force field ff14SB [43] was chosen for the protein. First, the domain was embedded in a truncated octahedral TIP3P [44] water box with a buffer of 15 Å, resulting in about 6690 water molecules. Neutralizing the system was achieved by introducing nine randomly placed Cl^-^ counterions, placed with the addion2 application, to minimize perturbations of the system by adding charges too close to the protein.

After minimization with harmonic restraints on protein-heavy atoms, the system was gradually heated from 100 to 300 K over 200 ps in NVT ensemble (see Wallnoefer et al. [45] for more details). A density equilibration over 1 ns was performed, followed by free simulations of the systems in NpT ensemble over 50 ns to ensure reasonable equilibration of the simulated systems. Production runs were carried out at 300 K using the Langevin thermostat [46] at 1.0 bar with an 8.0 Å nonbonded cutoff. A 2.0 fs timestep was chosen due to the usage of the SHAKE algorithm [47]. Snapshots were saved to trajectory every 500 steps or equivalent 1 ps for analysis.

Trajectories were analyzed using cpptraj from AmberTools16 [43] and PyMDLog (version 1.0.1). Distances and interactions of protein and ligand were likewise calculated via cpptraj. The resulting trajectories were analyzed for convergence of the usual MD simulation parameters (energies, density, temperature, etc.). Depictions were created with PyMOL (The PyMOL Molecular Graphics System, Version 2.3.2, Schrödinger, LLC).

### 2.3 Virtual Screening

#### 2.3.1 Structure-Based Virtual Screening

Selected frames were ranked accordingly to DrugScores [48]. The higher the score (which can be between 0 and 1) the more druggable the conformation is assumed to be [48]. A DrugScore above 0.6 and close to 1 corresponds to good druggability [48]. The protein conformations with the two highest DrugScores [48] were not investigated further as we found W134 dihedrals in these frames with low overall prevalence (See section 3.2.6). Therefore, we proceeded with the third highest-ranking TRMT2A RRM conformation (according to DrugScore [48]) and performed a virtual screening using MolPort stock compounds (7.409.777 compounds, www.molport.com) and NCI (237.771 compounds, https://www.nih.gov), retrieved as SMILES strings [49]. The libraries were prepared as follows: 1) With used built-in filters for lead-likeness [50] in the OpenEye tools (OpenEye Scientific Software, Santa Fe, NM. http://www.eyesopen.com. Hawkins, P.C.D.; Skillman, A.G.; Warren, G.L.; Ellingson, B.A.; Stahl, M.T.), 2) We discarded compounds in violation of the rule of five [51] and required strict atom typing (From the MolPort dataset 2.026.118 unique entries remained, for the NCI 49.657). Additionally, the compounds were adjusted to pH 7 and desalted. 3) A conformational expansion taking conformers of up to 15 kcal/mol into further consideration using OpenEye OMEGA [52] was performed. This resulted in 123 conformers on average per molecule for MolPort compounds and an average of approximately 42 conformers for NCI compounds. 4) The selected RRM domain MD snapshot was prepared with the apopdb2receptor application with W134 as the central amino acid 5) Next, a virtual screening with the outlined pocket definition using OpenEye FRED [53] with standard precision settings was performed. Conclusively, 1.188.185 MolPort, and 12.386 NCI compounds were successfully docked.

Finally, only compounds with a CHEMGAUSS04 better −7.5 were considered, and a subset of 13 compounds was selected based on a diverse set of non-covalent interactions with the RRM, visual inspection, availability, and costs for *in vitro* testing.

#### 2.3.2 Enrichment

A pharmacophore hypothesis based on compounds 13 and 15 was developed by using the “Small-Molecule Drug Discovery Suite” (Schrödinger Release 2018-2: Maestro). The 2D structures of compounds 13 and 15 were transformed into 3D structures by using the LigPrep application in Maestro (settings described in Supplement S2). The two compounds were selected, and the “Develop pharmacophore hypothesis” application was used to generate the pharmacophore model based on the selected structures (setting described in supplements S3). After ten pharmacophore models were generated, the highest-ranked model was manually edited. All features of the model were set to “required,” which means that every compound that matches the pharmacophore model must exhibit all features of the model. For testing the selectivity of the pharmacophore model in a screening, a decoy set for compounds 13 and 15 was generated with the webserver DUD.E (database of useful decoys: enhanced) [54]. None of the generated decoys matched the criteria of the pharmacophore model during a test screening. Therefore, the hypothesis was regarded as sufficiently selective.

The model was used to screen the MolPort database using the “Phase Ligand Screening” application in Maestro. The screening returned all molecules of the database with the respective screen scores. To eliminate structures with unfavorable properties, the scored structures were imported into Canvas (Schrödinger Release 2018-2: Canvas, Schrödinger, LLC, New York, NY, 2018). Only compounds with a screen score higher than 1,48 were retained. The remaining structures were converted into 3D structures, and molecular descriptors were calculated using the QikProp application. Compounds that were unlikely to possess ADMET properties favorable for CNS drugs were removed (details in supplements S4). Additionally, all molecules that did not fulfill the rule of five (No more than five hydrogen bond donors, no more than ten hydrogen bond acceptors, mass less than 500 daltons, logP not greater than five) or that did not pass the pan-assay interference compounds (PAINS) filter were eliminated. These are chemical compounds that often give false positive results in high-throughput screens [55]. A number of disruptive functional groups are indeed shared by many PAINS that tend to react nonspecifically with numerous biological targets rather than specifically against the selected target [56, 57].

To select ten representative structures of the remaining compounds, the structures were clustered and the ten most diverse cluster centroids were selected. To allow clustering, the structures were translated into binary vectors by using the molprint2D algorithm. The Tanimoto metric was used as a distance measure between the binary vectors. Based on a pairwise distance matrix, a hierarchical clustering with linkage type “McQuitty” was conducted. To determine the optimal number of clusters, the Kelley penalty score was calculated for each possible number of clusters. The set of clusters with the lowest score was considered as the optimal endpoint of the clustering.

The procedure described above was used to systematically sample compounds from the feature space constrained by compounds 13 and 15. Additionally, some compounds were sampled from a more focused feature space defined by highly similar derivatives of compound 13. Derivatives of compound 13 were returned by the MolPort search function after submitting a query with the IUPAC name of compound 13. Three very similar structures were selected based on visual inspection. The same explorative approach could not be used for compound 15 because the structure was no more commercialy available in MolPort.

#### 2.3.3 Ligand-Based Virtual Screening: Selection criteria for the pharmacophore

Some of the known TrmA inhibitors are based on the methylase activity requiring the cofactor S-adenosylmethionine (SAM or AdoMet) [58]. The SAM precursor L-methionine and structurally related norleucine are weak TrmA binders [59]. In addition, the ethyl analog of L-methionine (L-ethionine) is a known non-competitive selective inhibitor [59]. However, the related S-adenosyl-L-ethionine does not confer such selectivity [59]. On the other hand, ethylthioadenosine inhibits TrmA as well [59]. This suggests that both the adenosine and methionine substructures by themselves can mediate binding. Furthermore, several polyamines were shown to inhibit to a different extent *E. coli* TrmA, but highly dependent on environmental conditions [60]. While L-ethionine (15 mM) leads to 40% inhibition of tRNA methylation catalyzed by *E. coli* enzyme extracts it is, unfortunately, liver toxic in rats [59], [61]. The majority of the above-mentioned ligands are not toxic but fail to meet specificity requirements. Therefore, we decided to explore the chemical space around L-ethionine to circumvent toxicity-related issues, optimize the pharmacological/pharmacokinetics profile in general, while ideally retaining specificity. we chose, in addition, L-methionine and L-norleucine because they introduce a slight variability in the hydrophobic moiety.

First, low-energy conformers of the three selected known inhibitors were computed with ChemAxon tools (Calculator Plugins were used for structure, property prediction, and calculation, Marvin 19.8, 2019, ChemAxon (http://www.chemaxon.com)) of the three compounds and PharmaGist [62] to align them and build the hypothesis. This pharmacophore hypothesis was used to filter the ZINC database [63]. ADMET properties were computed with Canvas Version 2.8.014 (Schrödinger Release 2016-1: Canvas, Schrödinger, LLC, New York, NY, 2016.). Next, QikProp properties were calculated and all compounds violating the rule of 5 were discarded, while FDA-approved compounds were retained. Compounds were ranked according to human oral availability and blood-brain barrier permeability. After visual inspection, several known inhibitors from the literature and pharmacophore-derived compounds were selected for *in vitro* testing based on availability and associated costs.

### 2.4 Biophysical Experiments and Cell Assay

#### 2.4.1 Surface Plasmon Resonance (SPR)

SPR studies were performed using a BIACORE 3000 system (GE Healthcare). The TRMT2A RRM was diluted in 10 mM HEPES pH 7.5 for immobilization. Using coupling buffer (500 mM NaCl, 50 mM HEPES 7.5, 0.05% (v/v) Tween), the protein was amino coupled to a CM5 chip (GE-Healthcare) according to manufacturer’s instruction, reaching 3000 resonance units (RU).

Analysis of protein-compound interactions was performed at a flow rate of 30 μL/min in a running buffer (150 mM NaCl, 50 mM HEPES 7.5, 0.05% (v/v) Tween). The analyte of interest was diluted in a running buffer and injected for 2 min. To remove any residual attached compound, 2 × 2 min regeneration injections with 1 M NaCl were done in between runs. The concentration series reached from 0.5 μM to 8 μM for the positive control Spermine (Sigma Aldrich) and compound 12/Spermidine. For compounds 13, 15, 17, and 25, the concentration series reached from 0.5 μM to 32 μM.

Data were analyzed with the BIAevaluation software (GE Healthcare). The binding curves obtained were double-referenced against the signal in a protein-free reference channel and a buffer run. At the equilibrium of the binding curves, the corresponding response was plotted against analyte concentration. The KD was determined by fitting this curve to the steady-state affinity model. All experiments were performed in quadruplets on different days. Representative results for each compound are shown.

#### 2.4.2 Generation of HEKT293T with Stable RNAi-mediated Knockdown of TRMT2A

Stable RNAi-mediated knockdown of TRMT2A was achieved by infection of HEK293T cells with commercially available Lentiviral particles (MISSION^®^ shRNA Lentviral Transduction Particles NM_182984.2-**1574**s1c1; Sigma-Aldrich). Cells with stable integration of the shRNA construct were determined by a selection of puromycin-resistant colonies (0.5 μg/ml puromycin; Invitrogen). The efficacy of *TRMT2A* silencing effect was determined by qPCR and Western blot. The cell line with the strongest reduction of TRMT2A (>90%; short sh1574) was selected for use in further experiments. The same procedure was used to generate a control cell line (short shK) expressing a scrambled shRNA (SHC002V; Sigma-Aldrich).

#### 2.4.3 Transfections of HEKT293T cells

HEK293T cells were grown in Dulbecco‘s modified Eagle’s medium (DMEM) supplemented with 10% (v/v) FBS and 1% (v/v) penicillin/streptomycin. For transfection, cells were seeded at a density of 30 x 10^4^ cells per well in a 6-well-plate. Twenty-four hours post seeding, cells were transiently transfected with a construct mediating expression of an elongated polyQ with 103 glutamines fused to GFP (polyQ103:GFP) using either Metafectene^®^ (Biontex) or Calcium Chloride (CaCl_2_). Metafectene^®^ was used according to the manufacturer’s instructions, and CaCl_2_-transfection was achieved as follows: The transfection mix (in relation to the total volume of cell culture medium per well) contained 1% CaCl_2_ (1M), 4% ddH_2_O, 5% 2 x HBSS; 0.075% (v/w) DNA. The mix was incubated for 5 min and added to the medium of cultured cells.

#### 2.4.4 Inhibitor treatment

Inhibitors were added to fibroblasts (confluence of about 60%) by supplementation to the cell culture medium in a final concentration of 100 μM. In the case of HEK293T cells, inhibitors were subjected to cell culture media in a final concentration of 100 μM four hours post-transfection. Cell lysis was performed 48 hours after compound treatment for further analysis.

#### 2.4.5 Cell Death Assay

Treated cells were mechanically detached from the plate by scraping, and the cell/medium mixture was transferred to an Eppendorf tube. The cell/medium mixture was diluted in a 1:100 ratio with fresh DMEM medium. The diluted cell/medium mixture was supplemented with trypan blue solution (0.04%) in a 1:1 ratio. Cell viability was analyzed according to the dye exclusion test using the automated Cedex cell counter (Innovatis). For verification of observed differences in cell death, the effect of a few selected inhibitors was analyzed by a manual quantification of cell death. Here, trypan blue stained cells were analyzed in the Haematocytometer (Neubauer chamber). GFP-positive cells were counted, and the relative number of additional blue cells (dead cells) was determined. 100 GFP-positive cells were analyzed per condition, and three to six biological replicates were assayed.

#### 2.4.6 Cell lysis and protein fractionation

Cells were washed in PBS and then mechanically detached from the bottom of the plate in RIPA buffer (100 μl per six-well). Samples were centrifuged for 30 min at 13,300 rpm at 4 °C, and supernatants were collected (RIPA soluble fraction). The protein concentration of the RIPA soluble fraction was determined using the Bradford protein assay (DC Protein assay, Bio-Rad) according to the manufacturer’s instructions.

The pellet (RIPA insoluble fraction) was used for the Dot Blot analysis. The pellet (RIPA insoluble fraction) was washed twice (resuspended and centrifuged for 30 min at 13,300 rpm at 4 °C in a RIPA buffer). Finally, the pellet was resuspended in a urea buffer (30 mM Tris, 7 M Urea, 2 M thiourea, 4% CHAPS pH −8.5; 100 μl per six-well). Proteins in the pellet were solubilized by sonication (incubation in Ultrasonic Cleaner bath, VWR) for 10 min followed by incubation for 60 min at 4 °C under rotation. Subsequently, the lysate was centrifuged at 4 °C, 9,000 × g for 30 min. The supernatant was collected as the urea soluble fraction, and the urea insoluble pellet was discarded.

#### 2.4.7 Western blotting

20-50 μg of the RIPA soluble fraction protein samples were supplemented with 1x Laemmli buffer and boiled 95 °C, 5 min. Samples were subjected to SDS-PAGE (100 V for 90 min). Resolved proteins were transferred onto nitrocellulose membrane (225 mA per gel for 60 min) by semi-dry blotting. The membranes were then blocked with skimmed milk (5% in TBS-T for 60 min) followed by incubation with the primary antibody at 4°C overnight (in 5% skimmed milk in TBS-T). The primary antibodies used were mouse anti-Polyglutamine, Milipore MAB1574; mouse anti-Huntingtin polyQ, Developmental Hybridoma Bank, MW1 or mouse anti-GFP; Roche 11814460001. All antibodies were applied in a 1:1000 dilution. Membranes were washed three times for 10 min in TBS-T and incubated with the secondary horse radish peroxidase (HRP) coupled antibody (GE Healthcare, NXA931V; in a 1:3000 dilution) for two hours under agitation at room temperature. Subsequently, the membranes were washed three times in TBS-T for 10 min, and the chemiluminescence signal was detected using the Super Signal^®^ West Femto Maximum Sensitivity Substrate (Thermo Scientific) detection kit according to manufacturer’s instructions. Chemiluminescence signals were visualized using the Alliance UVltec system (Biometra) and processed using Image J software.

#### 2.4.8 Filter retardation assay

Relative to the total protein concentration in the RIPA soluble fraction (volumes kept constant), we loaded a volume that would correspond to 90 μg of protein in the RIPA soluble fraction. The total volume was adjusted to 50 μl by addition of RIPA buffer supplemented with a 7 μl dot blot buffer (0.5 M Tris pH-6.8, 0.4% (w/v) SDS, 20% (v/v) glycerol, 0.2 M DTT). The samples were transferred to a nitrocellulose membrane (Protran^®^BA 83, Whatman, 0.2 μM pore size) by sucking the probes through the membrane using a vacuum pump attached to a dot blot chamber (Camlab, UK). The membrane was washed several times with TBS-T and blocked for 1 hour using 5% skimmed milk in TBS-T. Afterward, the membrane was treated as described for Western blotting using respective primary and secondary antibodies.

#### 2.4.9 Reverse-Transcription Quantitative PCR (RT-qPCR)

To assess the mRNA levels of TRMT2A in shK and sh1574 cells, we utilized RT-qPCRs. After dissolving and homogenizing cells in PeqGold (PeqLab #30-2010, Germany), total RNA was extracted using peqGold RNA TriFast as previously described [64, 65]. Complementary DNA was synthesized using the MMLV reverse transcription kit (Cat.# 28025-013, Thermo Fischer Scientific, Waltham, Massachusetts, USA) and random hexanucleotide primers (Cat.# 48190-01, Thermo Fischer Scientific, Waltham, Massachusetts, USA) using 1 μg of total RNA. Triplicates of every sample were transferred by a pipetting robot (Corbett CAS-1200, Qiagen, Hilden, Germany) to Rotor-Gene strip reaction tubes (Starlab, 22143, Hamburg, Germany) and RT-qPCR analysis was performed using the Rotor-Gene Q device (Qiagen, 40724, Hilden, Germany). RNase free H2O (Merck-Millipore, 64293, Darmstadt, Germany) served as no template control (NTC) and primer efficiencies were calculated using the Pfaffl method [66]. The target gene *TRMT2A* and the housekeeping gene hypoxanthine guanine phosphoribosyltransferase (*HPRT)* were measured at cycle threshold (Ct values) and relative quantification was calculated by the ΔΔCt method method using the qbase+ software (Biogazelle, Belgium). Data are expressed as relative amount of the target genes to *HPRT*. The following forward (fwd) and reverse (rev) primers were used ( ): *TRMT2* (fwd: CTG CAG AGC CCC ATC TAA CC; rev: TTA TCT GGG GGT CCT GGT GT), *HPRT* (fwd: ACC CCA CGA AGT GTT GGA TA; rev: AAG CAG ATG GCC ACA GAA CT).

#### 2.4.10 Statistics

GraphPad Prism 5.0 software was used for statistical analysis and data presentation. Mean and standard deviation (SD) is presented in all graphs. Tests used for comparisons of individual bars in graphs are noted in the figure legend. p < 0.05 was considered significant. In graphs, p values are depicted as *p < 0.05; **p < 0.01; ***p < 0.001, ns = not significant.

### 2.5 LC-ESI-MS analysis

Prior to measurement, the samples were centrifuged at 20,800 g and 21 °C for 10 min, followed by a filtration through an *AcroPrep Advance 96well Supor plate* (0.2 μm) into a 96well PCR plate by centrifuging 40 min at 3220 g and 21 °C. The LC-ESI-MS analysis of the samples - containing both the natural nucleoside and 13 pmol of heavy-atom labeled ^13^C_5_-m^5^U as internal standard – was performed on a *Dionex Ultimate 3000* HPLC system coupled to a *Thermo Fisher LTQ Orbitrap XL* mass spectrometer. For the chromatography, an *Interchim Uptisphere120-3HDO C18* column was used whose temperature was maintained at 30 °C. Eluting buffers were buffer A (2 mM HCOONH_4_ in H_2_O (pH 5.5)) and buffer B (2 mM HCOONH_4_ in H_2_O/MeCN 20/80 v/v (pH 5.5)) with a flow rate of 0.15 mL/min. The gradient was: 0→3.5 min, 0% B; 3.5→4 min, 0→0.2% B; 4→10 min, 0.2% B; 10→70 min, 0.2→7% B; 70→100 min, 7→60% B; 100→102 min, 60→100% B. The elution of the nucleosides was monitored at 260 nm with a *Dionex Ultimate 3000 Diode Array Detector*, and the chromatographic eluent was directly injected into the ion source of the mass spectrometer without prior splitting. Ions were scanned in the positive polarity mode over a full-scan range of *m/z* 225-600 with a resolution of 100,000. Parameters of the mass spectrometer were tuned with a freshly mixed solution of adenosine (5 μM) in buffer A and set as follows: Capillary temperature 275 °C; capillary voltage 20 V; source voltage 3.50 kV; tube lens voltage 55 V. The relative m^5^U-content of the samples was determined using the program *Qualbrowser*: For each measurement, the ion chromatograms of natural m^5^U and its corresponding heavy-atom labeled isotopologue were extracted from the total ion current (TIC) chromatogram, and the ratio of the peak areas was calculated. Each sample was measured as experimental duplicate.

## 3. Results and Discussion

Silencing the *Drosophila* homolog of TRMT2A strongly suppressed polyQ-induced toxicity and polyQ aggregation in flies [12]. As TRMT2A is well conserved, we asked whether pharmacological inhibition of human TRMT2A would also exert effects on polyQ-affected proteins as observed in flies.

To develop inhibitors against human TRMT2A, two parallel strategies were pursued. On the one hand, the structural and dynamic properties of the N-terminal RNA recognition motif (RRM) of TRMT2A were analyzed by solving the RRM crystal structure and leveraging this information with computational approaches. The TRMT2A RRM structure was analyzed to identify putative druggable binding pockets and subsequently enabled a structure-based virtual screening campaign. On the other hand, a ligand-based pharmacophore model across known tRNA methyltransferase inhibitors was built to find possible molecules targeting the methyltransferase catalytic domain (CD) of TRMT2A. The set of ligands identified from these two procedures was tested in cellular experiments. A selection of those targeting the TRMT2A RRM was also tested in biophysical experiments.

### 3.1 Structure Determination of the TRMT2A RNA Recognition Motif

A variety of sub-fragments of TRMT2A were designed, cloned, and expressed in *E. coli*. Amongst them, only a fragment from amino acids 69 to 147, which essentially included the RRM, could be expressed and purified to near homogeneity. Robot-assisted crystallization screens yielded initial hits that were subsequently optimized for larger, well-diffracting crystals. An initial structure was solved from a dataset collected to 2.0 Å resolution (R_free_ = 25.14%, German Electron Synchrotron DESY; Table S1), and an improved structure was determined at 1.23 Å resolution with favorable statistics and stereochemistry (R_free_ = 17.15%, Swiss Light Source (SLS); Table S1). The structures revealed the typical arrangement of four anti-parallel β-strands and two α-helices that are packed against each other. They are arranged in a β1-α1-β2-β3-α2-β4 fold topology (Figure 1A), as already reported for other RRMs [68]. The root mean square deviation of its Cα atoms to its closest structural homolog (PDB-ID: 3vf0 [69]) is 1.5 Å and 1.3 Å in the 2.0 Å and 1.2 Å structure, respectively. Homology search has been performed with DALI server [70]. Loops that connect β1/α1 (referred to as loop 1), β2/β3 (loop 3), and α2/β4 (loop5) are located on the side featuring the β-sheet, whereas loop 2 (α1/β2) and 4 (β3/ α2) are closer to the α-helices as outlined in Figure 1A. Typically, the β-sheet of an RRM interacts with a variable number of two to eight unpaired RNA nucleotides via two consensus regions located in the center of the two neighboring β-strands. These sub-motifs are referred to as ribonucleoprotein (RNP) 1 and 2 [71]. In the structure of this RRM, known determinants of RNA-RRM molecular recognition can be distinguished, such as positively charged residues and aromatic amino acids, which form π-stacking interactions with the RNA nucleobases [72].

**Figure 1.**
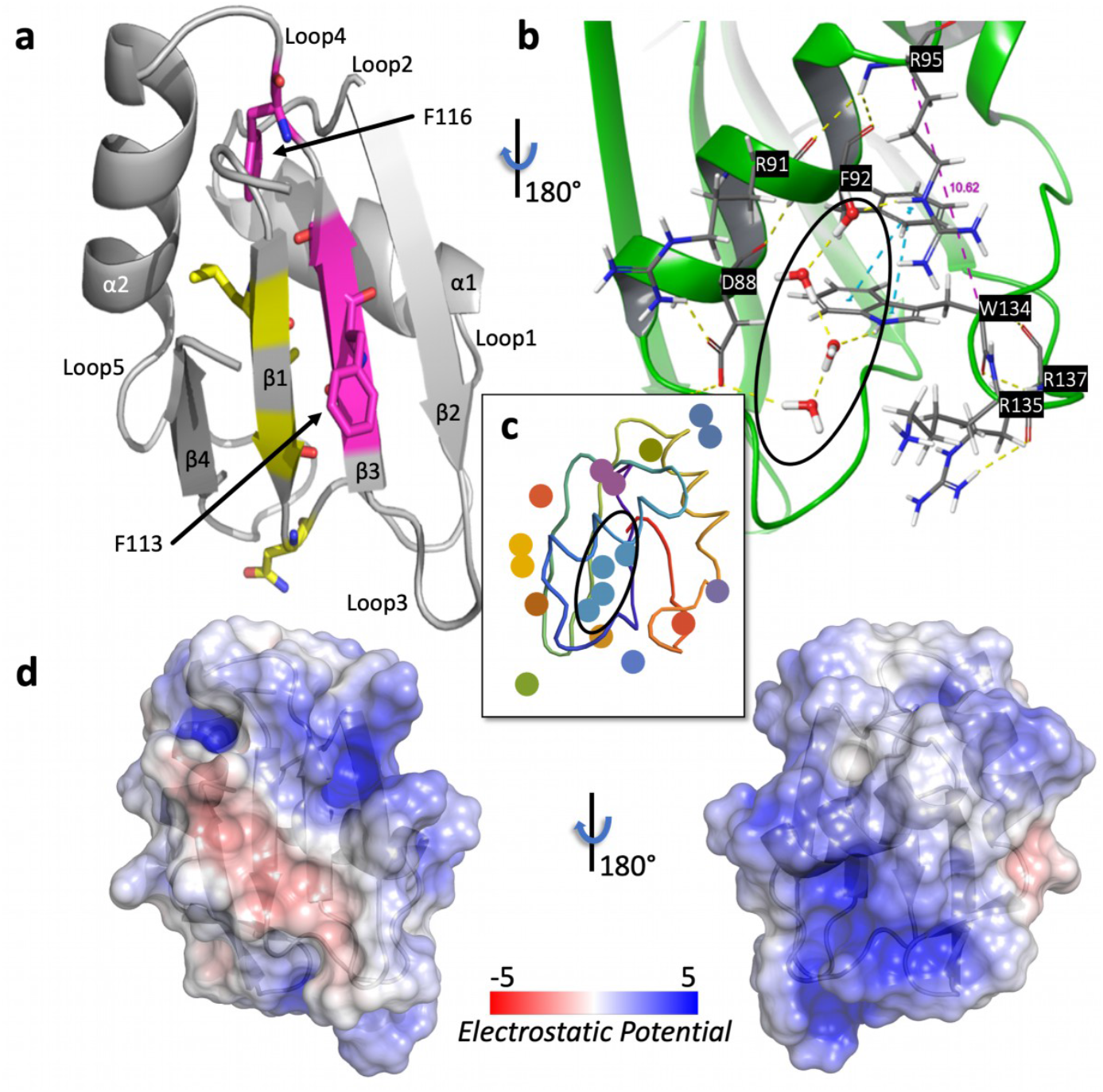
Structural features of TRMT2A RRM (resolution: 2.0 Å). (A), X-ray structure of the RRM in cartoon representation. The two central β-strands feature the submotifs RNP1 (β3 strand) and RNP2 (β1 strand). Canonical RNP1/2 residues found in TRMT2A RRM are highlighted (RNP2 as yellow sticks on the β1-strand and RNP1 as magenta sticks on the β3-strand). The sidechain of F113, part of RNP1, is solvent-exposed. In contrast, the sidechain of F116 (part of RNP1 as well) is buried and part of loop 4. (B), View of the lower helical site of the RRM and the water network. This view of the protein was obtained by rotating the protein’s orientation shown in A by 180 degree. The sidechain of W134 π-stacks with F92 (blue dashed lines). The Cα distance between R95 and W134 is highlighted as a purple dashed line (see subsequent sections). (C), Agglomerative Euclidean-distance-based clustering. Discs with the same color belong to the same water cluster. The largest cluster is circled as in B. (D), Electrostatic surface obtained with the Adaptive Poisson-Boltzmann Solver (APBS) [67] Electrostatics PyMOL plugin and mapped onto the solvent accessible surface in units of kT/e. The same protein’s orientations in A and B, respectively, were preserved.

Here, RNP1 was shown to be composed of K/R-G-F/Y-G/**A112**-**F113**/Y-**V114**/I/L-X-**F116**/Y (boldface amino acids refer to positions found in TRMT2A RNP1/RNP2; X refers to an arbitrary amino acid) [73]. RNP1 is present in the RRM β3 strand of TRMT2A and spans from aa 109 to 116. RNP2 features I/V/**L75**-F/Y-I/V/**L77**-X-**N79**-L; RRM RNP2 is part of the adjacent (aa 75-80) β1 strand. RRMs often display two conserved aromatic residues on RNP1 and RNP2, typically phenylalanine or tyrosine residues. One aromatic RRM residue is frequently π-stacking with the 3’ end of the RNA, and another is inserted between two RNA ribose rings [72]. The positioning of the aromatic side chains is crucial for RNA recognition [74]. In our structures of the TRMT2A RRM, two phenylalanines are present on RNP1, but only one exposes its sidechain to the solvent (F113).

### 3.2 Structure-based virtual screening on RNA Recognition Motif of TRMT2A

Neither of our determined crystal structures of TRMT2A RRM reveals distinct binding cavities (e.g., starting points for structure-based ligand discovery). Therefore, traditional druggability detection methods were augmented with alternative approaches, which incorporate protein conservation, solvation, and local flexibility considerations. Using these approaches, a potential cryptic site was uncovered. Cryptic sites are not evident in the unbound protein surface, but they can extend into underlying cavities upon ligand binding, providing tractable drug target sites and thus expanding the druggable proteome considerably [75].

#### 3.2.1 Bioinformatics Analysis

Binding hotspots have been relatively well conserved during evolution, unlike the remainder of the protein surface [76]. Analysis of conservation patterns in the amino acid sequence is, therefore, a useful tool to detect binding sites.

To this aim, RRMdb, an evolutionary-oriented database for RNA recognition motif sequences [77] was queried, where the RRM of TRMT2A is classified as belonging to the family 262 with 25 known members from eukaryotes. The sequence conservation of the RRM residues was calculated via the multiple sequence alignment of these 25 sequences (see Methods and Figure S1 A-B for details). Highly conserved residues are not only found on the face of the β-sheet (the canonical RNA binding site) but also on the helical back (see Figure 1 B and S1 C). All aligned sequences contain a lysine at position 135 (human TRMT2A numbering). A tryptophan at position 134 is present in all but three of the 25 sequences (88% of conservation), where an aromatic residue, tyrosine, or phenylalanine takes its place. Interestingly, R95 is also highly conserved (77%). Such high sequence conservation, far from the putative tRNA binding site, suggests a functionally important role of these residues. Specifically, tryptophan, given its bulky and rather hydrophobic sidechain, is indeed rarely found in loops, unless these loops are involved in protein-protein interactions [78].

#### 3.2.2 Crystallographic Water Clustering and 3D-RISM Analysis

Since water clusters can mediate protein-ligand binding, these were searched at the surface of the protein [79]. An agglomeration-based clustering of water molecules in the 2.0 Å crystal structure was performed, and four non-singleton clusters were identified (see Figure 1B-C). The largest crystallographic cluster contains four water molecules and is located between α1 and loop 5 in the so-called *minor/small groove*. In the 1.23 Å structure, a sulfate ion takes the place of this cluster.

This motivated us to investigate solvation effects in more detail with the three-dimensional reference interaction site model (3D-RISM) theory [80]. The latter can estimate water distribution propensities of each solvent site within and around a solute (see Figure S2). This analysis confirmed the hydration site centers we found in the X-ray structure with 2.0 Å resolution. Notably, two ‘unfavorable’ solvent sites buried 2.5 Å beneath the sidechain of W134 were detected as well: the favorable release of corresponding water molecules to the bulk may drive local structural rearrangements.

#### 3.2.3 Drug Binding Site Prediction on the Protein Crystal Structure

Next, a druggability assessment was performed, i.e. the size, shape, and physicochemical features of all RRM cavities were analyzed with DoGSiteScorer [38] (v2.0.0_15.01.2019). Five candidate binding sites were identified (see Figure S3) in the 2.0 Å structure, amongst which the largest one coincides with the above-mentioned *minor/small groove*. A corresponding but equally inaccessible site was also identified in the 1.23 Å structure (see Figure S4A). All other sites detected herein were found to be too shallow to accommodate a potential small molecule ligand.

The possibility of cryptic site formation was therefore investigated. Cryptic or transient sites are defined as only opening up when a binder is present [75]. In a recent attempt to expand the druggable proteome, these sites were suggested as an interesting alternative for hit discovery, particularly for proteins lacking conventional binding pockets [39].

#### 3.2.4 Transient Site Discovery on TRMT2A

CryptoSite [39], which allows the rapid identification of residues contributing to cryptic site formation via a support vector machine (SVM), was used. We found that the highly conserved residues W134 and K135 are likely part of a cryptic site located in the *minor/small groove* (Figure 2A). Further analysis of this site with implicit solvent molecular dynamics simulations conducted with TRAPP (TRAnsient Pockets in Proteins [40]) corroborated this finding. Specifically, below the *minor/small groove*, formed in part by W134/R135, an accessible cavity is repeatedly detected upon different models of perturbation: either under rotamerically induced site chain perturbations with Langevin dynamics (L-RIP) [41] or via ligand-induced perturbations (RIPLig) with a probe mimicking an apolar/aromatic ligand approaching the *minor/small groove* region (Figure 2B-D and for the 1.23 Å structure Figure S4B) [41].

**Figure 2.**
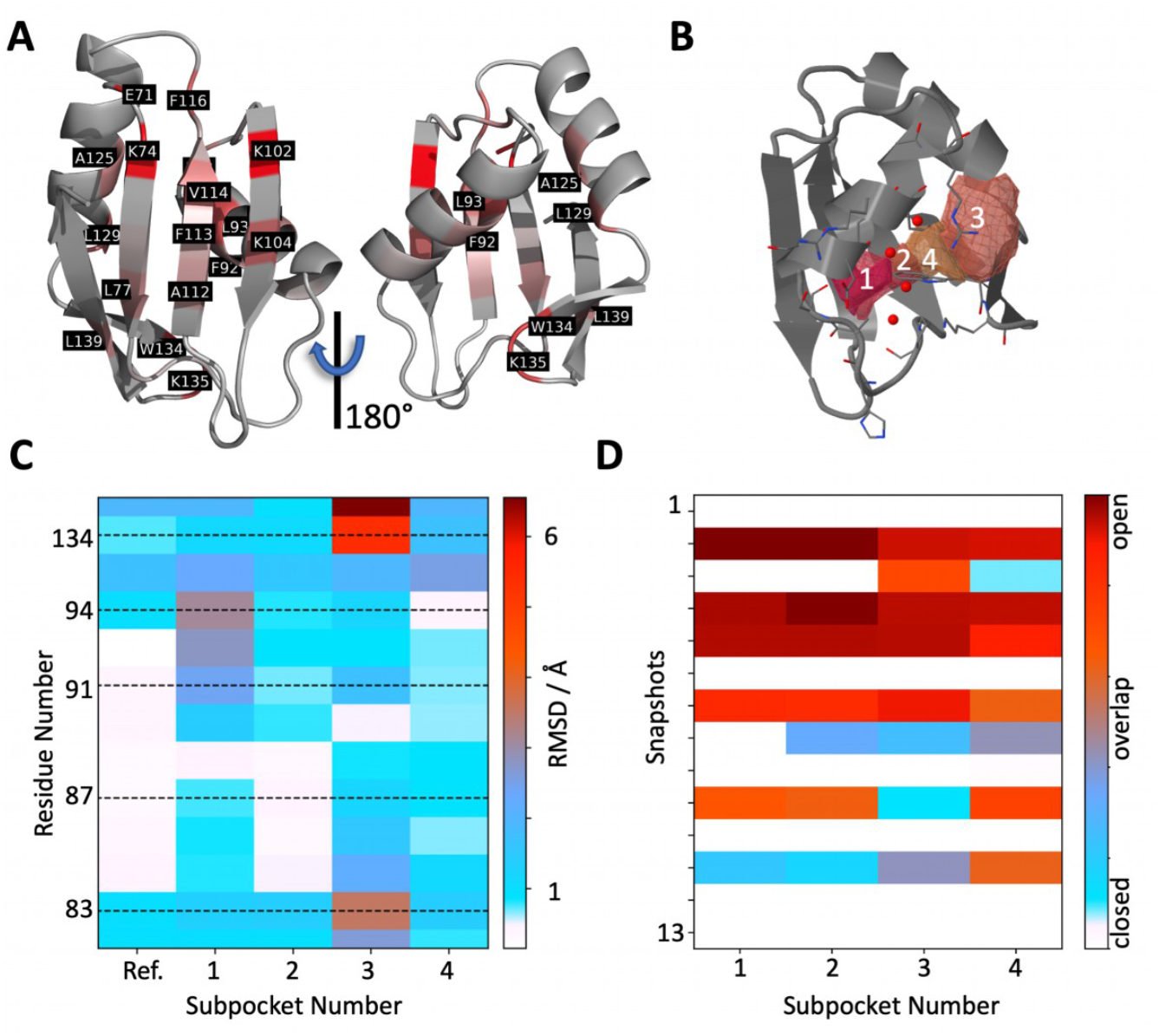
Cryptic site assessment. (A), CryptoSite indices mapped onto the TRMT2A RRM (shown in cartoon representation). Values ranging from gray, unlikely part of a cryptic site to red, higher probability of contributing to a cryptic site. On the helical site of the RRM, W134 and L135 were detected as putative contributors to cryptic site formation. (B), Rear view of the RRM with the identified subpockets and consistent numbering in b, c, and d with TRAPP. All appearing transient regions are depicted as isosurfaces (at 25% occurrence during the simulation). Binding site residues are depicted as sticks. (C), Representative TRAPP binding site flexibility estimation (RMSD values in Å are color-coded from white to red) using the L-RIP/RIPlig approaches. W134, H83, and K135 show high structural variance. (D), Subpocket occurrence during the simulations shows that all four subpockets are formed repeatedly.

Our findings are in line with the fact that the *minor/small groove* of other RRMs has been previously reported to interact with different peptides [72]. However, our predicted transient site formation in the *minor/small groove* region is rather distant from the canonical RNA binding site and might not necessarily inhibit RNA binding. Therefore, this transient site needs to be allosterically connected to the canonical tRNA binding site, for any binders in the minor/small groove to be able to modulate tRNA binding. To support such hypothesis, the possibility of allosteric communication occurring between this identified transient site and the putative tRNA binding site (RNP1/2) was investigated.

#### 3.2.5 RNP1/RNP2 and small groove allosteric regulation

Perturbations mimicking ligand binding on the suggested tRNA-anchor-phenylalanine (F113 of RNP1) were simulated to see if it can cause a functional change in the *minor/small groove* region and thus predict the presence of allosteric communication. For this purpose, both structures were analyzed with AlloPred [42], which uses perturbation of normal modes alongside other pocket descriptors in a machine learning approach. These results indicate that the *minor/small groove* region can allosterically modulate the β3-strand where F113 is present (see Figure S5 for the 2.0 Å structure and Figure S4C for the 1.23 Å structure, respectively). Specifically, in our analysis, W134 seems to be a major player in this allosteric communication since changes in its immediate surroundings are allosterically connected to the suggested tRNA binding site. This is in line with our water propensity distribution and conservation analysis, pointing towards a key role of this residue.

#### 3.2.6 Molecular dynamics and snapshot selection

To corroborate our findings, the flexibility of this putative allosteric pocket was investigated through molecular dynamics (MD) simulations. A 10 μs-long MD trajectory in explicit solvent was performed (Figure S6). While the overall structure remained stable, loop 5 appears to contribute most to the flexibility of the RRM. This is in line with the water distribution analyses in this area discussed above. We found that the C_α_ distance of W134 and R95 can increase from 10.62 Å in the crystal structure to more than 20 Å (Figure 3A) during the MD simulation, as can be seen in a representative MD snapshot (Figure S7). Indeed, W134 transiently passes from buried conformations, where π-stacking with F92 is possible (Figure 1A and 3B), to solvent-exposed ones. The conformations where W134 is solvent-exposed are also characterized by loop 5 structural rearrangements (further details are provided in Figure S8 A-B of the SI). When the C_α_ atoms of conserved amino acids R95 and W134 are in close proximity, the consistent formation of an enclosed cavity during the MD simulation of the 2.0 Å structure of the RRM was observed. Moreover, an opening in the 1.23 Å structure (Figure S4 D) was also detected. Here, the indole ring of W134 is gradually pushed outwards and becomes solvent-exposed. This is of particular interest since W134 was indicated in our initial CryptoSite model, as well as in the follow-up TRAPP analysis, as key residue involved in putative cryptic site formation, together with R95.

**Figure 3.**
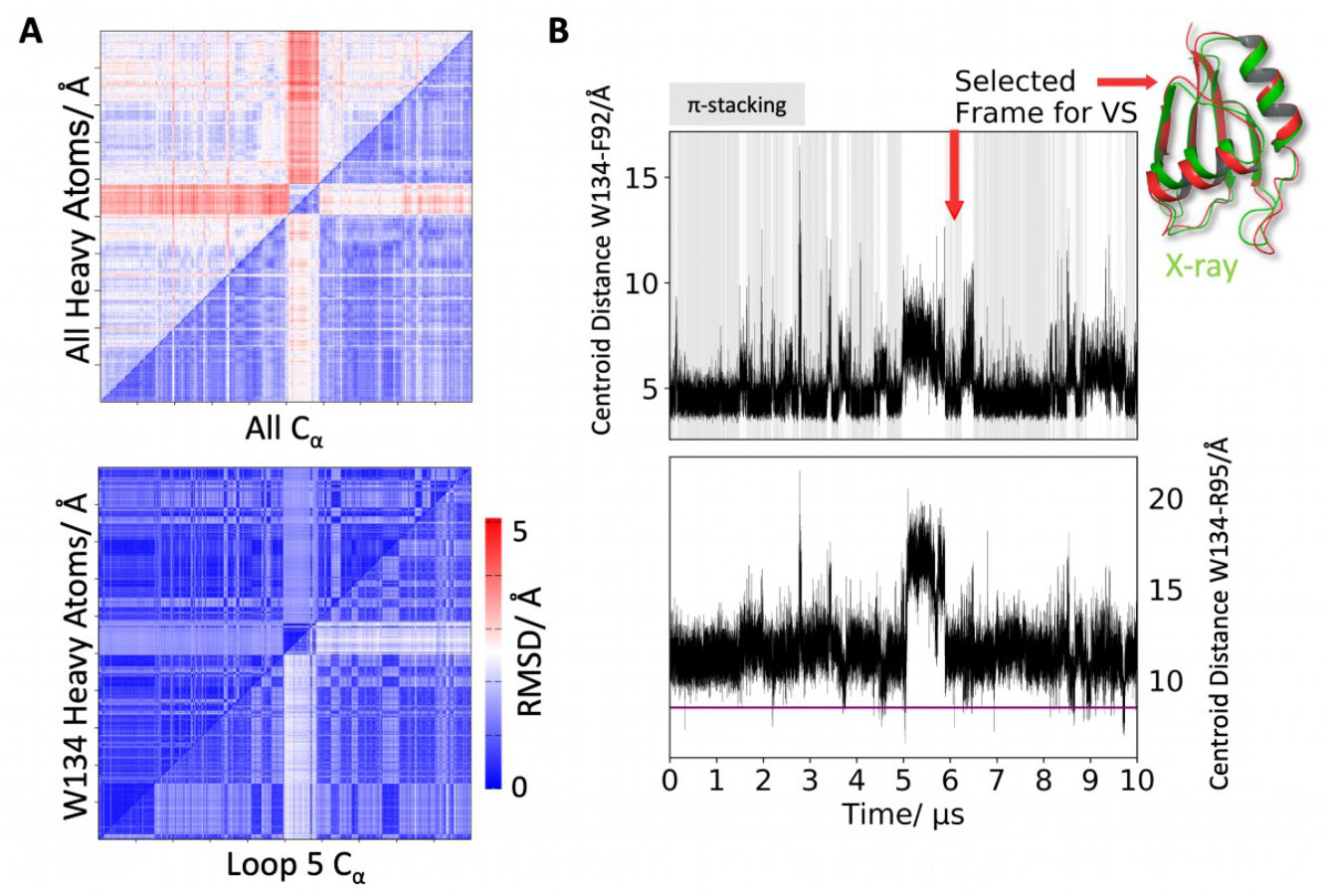
Explicit-solvent molecular dynamics results. (A) Combined2D-RMSD plots of Cα atoms (horizontal axes) and heavy atoms (vertical axes) over the 10 μs-long simulation. (top) Heavy atoms/C*α* RMSDs indicate an overall stable structure, with considerable rearrangements occurring after about 5 μs. (bottom) Flexibility in the loop 5 region is mediated by the transient π-stacking of W134 and F92 (blue dashed lines in Figure 1B); (B) (top) These structural changes are also reflected in the centroid distances of the aromatic rings involved. Frames with detected π-stacking, according to reference *[81]*, are underlined in gray. (bottom) Cα distance between R95 (α1 helix) and W134. We consistently observed the formation of a hydrophobic pocket when these residues approach. Thus, we extracted snapshots with an R95-W134 Cα distance below 8.5 Å (indicated by the magenta line) for a detailed druggability analysis. The red arrow indicates the snapshot (red) used for virtual screening (VS) and is superimposed with the low-resolution crystal structure (green).

Thus, frames with a C_α_ distance (R95-W134) cut-off of 8.5 Å were selected for further analysis (104 frames out of 100,000 total frames for the whole trajectory). Among these, six conformations had a DrugScore [48] (see methods) >0.60. The two highest-ranking receptor conformations displayed a W134 indole ring flip, which occurs infrequently during the simulation (See Figure S8 C), and were therefore not considered. The third highest-ranking conformation, however, was frame 60,942 (after 6,094.2 ns, DrugScore [48]: 0.62), where the loop is in the process of refolding to its crystallographic conformation after a roughly 1 μs long reorganization period. In this state, the volume of the *minor/small groove* is significantly increased, reaching up to 589 Å³ (See Figure S8 D).

By comparing this selected conformation with the original X-ray structure, we can see that the loop α2-β4 moves outward (Figure 3B). During this widening of the pocket, the side chain of tryptophan (W134) opens a small and transient cavity below the small groove. The indole ring of W134 then forms a cation-*π* interaction with R135. This interaction can energetically compensate for the unfavorable solvent exposure of the indole ring and breaking the *π*-stacking with F92. The RMSD between the crystal structure and the selected structure is 1.74 Å (Figure 3B).

This partially hydrophobic pocket was next used for structure-based small inhibitor discovery. A more detailed pocket description was elucidated with Schrödinger SiteMap and is shown in Figure S8 E [82]. In addition, Replica Exchange simulations (REST2) [83] with solute tempering were performed (see Supplement S1, Figure S9, and Figure S10) to enhance the exploration of biomolecules’ conformational space [84]. Our findings suggest that the opening of the cryptic pocket is a rare event that might be facilitated by ligand binding.

#### 3.2.7 Virtual Screening on TRMT2A RRM and Post Processing

To exploit this putative cryptic pocket, a structure-based virtual screening workflow was performed to prioritize small molecules that might bind into this site for *in vitro* testing. Virtual screening can be defined as a set of computational methods that analyzes large databases or collections of compounds to identify potential hit candidates [85]. As a first step, therefore, *in silico* collections of commercially available small molecules were retrieved and prepared as outlined in the methods section. Next, the curated compound library was docked into the identified binding site. A possible binding pose for each ligand conformer was generated and then prioritized with a fast-scoring function (see methods). Ligands with unreasonable binding mode hypotheses were discarded, while the best ones were retained and subsequently rescored using a more detailed scoring function (see methods). Among these, 13 compounds were selected by visual inspection to undergo experimental testing (compounds 13-25, Table S2). Further selection criteria were based on the diversity of interactions formed with the TRMT2A RRM (Figure 4 A) and availability. Notably, several compounds predicted to bind into the *minor/small groove* revealed a common substructure (Figure 4 A-B), despite displaying otherwise diverse binding modes. Therefore, we decided to enrich our original pool with the above-mentioned substructure and performed based on a pharmacophore (i.e. an abstract description of the molecular features that are believed to be necessary for ligands to be biologically active against the selected target protein) obtained from compounds 13 and 15. Three compounds were added as results of such a procedure (see methods for details).

**Figure 4.**
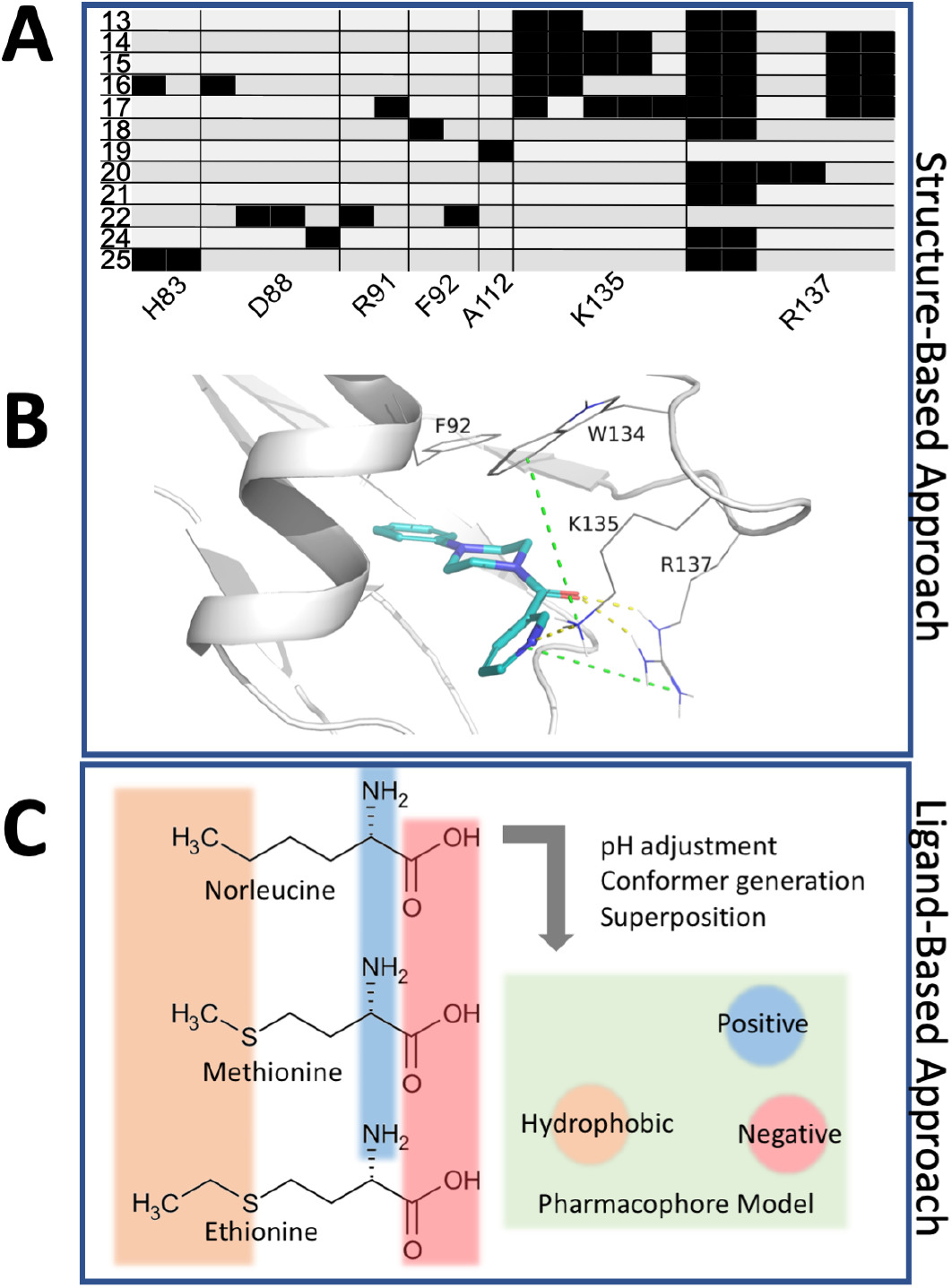
Predicted protein-ligand interaction patterns. (A), Protein-Ligand Interaction Fingerprint (PLIF) of docking poses into the cryptic site depicted as barcodes. Black bars indicate one or more interactions between TRMT2A RRM residues (columns) and individual ligands (rows). The first column indicates the running number of the docked compounds. (B), Interactions of the docked pose of compound 13. K135 and R137 form hydrogen bonds with the ligand. Moreover, the phenyl ring of the ligand can potentially π-stack with F92. Cation-π interactions between K135 and W134, but also between R137 and the pyridinyl ring, are apparent (green dotes lines). (C), Pharmacophore for three structurally similar compounds that inhibit the E. coli homolog of TRMT2A. The angle between positive-negative-hydrophobic features is nearly a right angle (86.9°). The distance between the hydrophobic and the positive/negative (ionic) feature are 7.3 Å and 6.8 Å, respectively.

### 3.3 Ligand-based virtual screening on the methyltransferase catalytic domain of TRMT2A

To identify TRMT2A inhibitors binding to the CD, a ligand-based virtual screening was performed. The latter uses the information present in known active ligands rather than the structure of the target protein. This is a suitable approach here since we lack structural information on the TRMT2A CD. Specifically, another pharmacophore hypothesis was generated and exploited to screen libraries of chemical compounds to elucidate those with the required molecular and spatial features for binding to this site.

In this ligand-based virtual screening, we searched for all available inhibitors of TRMT2A homologs. Unfortunately, only data for TrmA in *E. coli* were found [58-61]. Therefore, the known TrmA inhibitors L-norleucine, L-methionine, and L-ethionine were superimposed, and a pharmacophore model was built. The selection criteria of these three are explained in detail in the methods. The resulting pharmacophore model contained only three features: a hydrogen bond donor and acceptor (amino acid abstraction) and an approximately equidistant hydrophobic site feature, representing apolar sidechains (Figure 4C). Next, this pharmacophore was used to screen libraries of known chemical compounds to identify the ones with the same selected molecular features. Compounds were sorted according to predicted absorption, distribution, metabolism, and excretion (ADMET) properties, human oral availability, and blood-brain barrier permeability. After visual inspection, several known inhibitors from the literature and pharmacophore derived compounds were selected for experimental testing (compounds 1-12, Table S2).

### 3.4 *In cell* assays

Human embryonic kidney cells (HEK293T) were used to determine the effect of *TRMT2A*-silencing on polyQ-induced toxicity and aggregation. For analysis, we first generated cells with stable expression of a short hairpin RNA (shRNA) to induce *TRMT2A* specific RNAi (sh1574). A cell line with stable expression of a scrambled shRNA served as control (shK). Analyzing the abundance of TRMT2A levels in both cell lines revealed a strong reduction of TRMT2A abundance/protein levels compared to shK cells in Western blot analysis (Figure 5 A, B). Similarely, qRT-PCR analysis revealed a significant reduction of *Trmt2a* transcript abundance in sh1574 cells compared to shK cells (Figure 5 C). In the next step, we asked whether reduced TRMT2A abundance decreased polyQ aggregation as observed in flies [12]. We transiently transfected sh1574 and shK cells with plasmids coding *huntingtin* exon1 harboring a polyQ stretch of 25 or 103 glutamines fused to GFP (HttQ25ex1:GFP or HttQ103ex1:GFP, respectively). No differences were observed in the abundance of soluble HttQ25ex1:GFP or HttQ103ex1:GFP in lysates derived from shK cells compared to sh1574 cells. PolyQ aggregates were analyzed by filter retardation assay (FRA) two days post-transfection [86]. The FRA analysis revealed a strong aggregate load in shK cells. However, in sh1574 cells, the abundance of polyQ aggregates was low (Figure 5 D).

**Figure 5.**
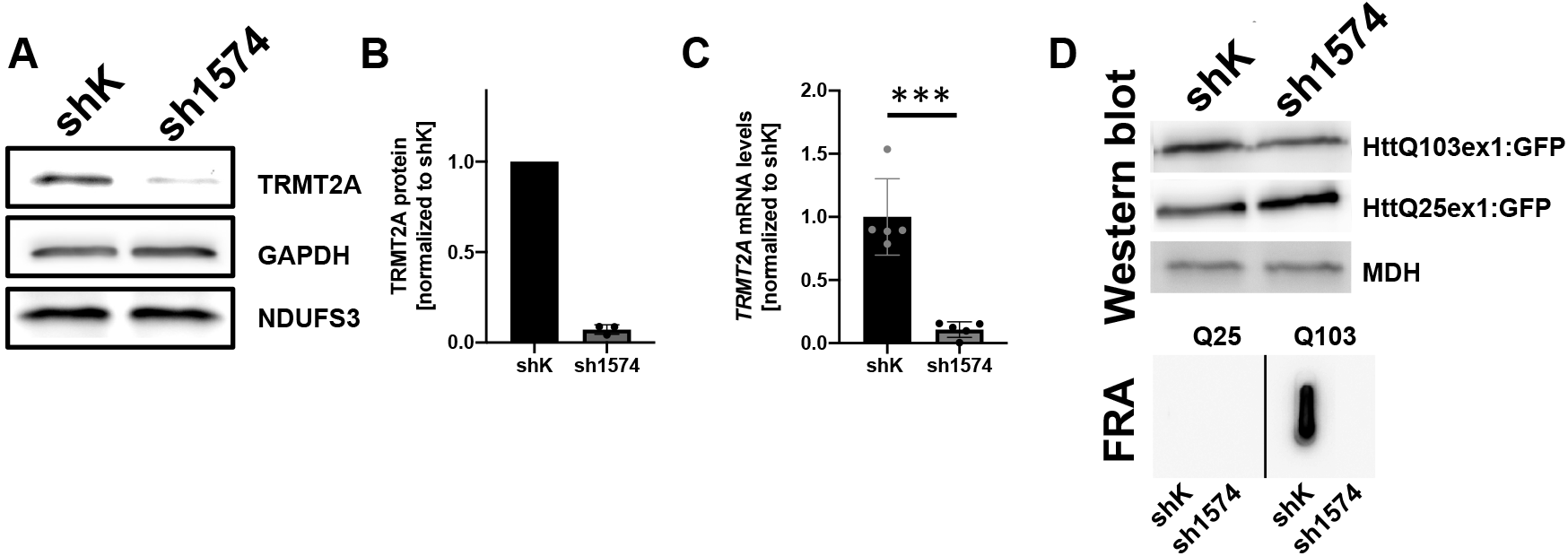
TRMT2A-deficient cells display reduced PolyQ aggregation. (A) Representative Western blot analysis to detect TRMT2A abundance in cell lysates derived from control (shK) and stable TRMT2A-silenced cells (sh1574). Of note cell lysates were generated from 1 million cells each and not adjusted on protein concentration. Detection of GAPDH and NDUFS3 were used for normalization. (B) Bar graph depicting differences in TRMT2A abundance in shK and sh1574 cell lysates determined by Western blot quantification (N=3). (C) Bar graph depicting differences in mRNA abundance of *Trmt2a* comparing shk and sh1574 cells. Ordinary two-tailed t-test was performed to determine significance (***p<0.001). (D) Detection of exogenous HttQ103ex1:GFP and HttQ25ex1:GFP peptides by Western blot analysis after transient transfection of shK and sh1574 cells. No difference in the abundance of GFP-tagged peptides was detected in the RIPA soluble fraction. Detection of Malate Dehydrogenase (MDH) was used as the loading control. RIPA insoluble HttQ103ex1:GFP and HttQ25ex1:GFP aggregates were analyzed by Filter retardation assay (FRA). No insoluble aggregates where detected for HttQ25ex1:GFP, HttQ103ex1:GFP aggregates were present in lysates derived from shK cells, but strongly reduced/absent in lysates derived from sh1574 cells.

In addition, we tested sh1574 for alterations in overall protein synthesis, cell viability and cell growth. Compared to shK cells, sh1574 did not show any obvious difference in the anlyzed parameters (Figure S11). Having established both cell lines, we used them to address whether an inhibition of TRMT2A might cause a reduction in polyQ aggregation and polyQ-induced toxicity. Indeed, while, expression of HttQ103ex1:GFP in shK cells resulted in robust polyQ aggregate formation, sh1574 cells displayed strongly reduced polyQ-aggregates (Figure 5 D, Figure 6 A). Assuming a specific inhibition of TRMT2A with selected compounds, we would expect a strong decrease in polyQ aggregate abundance. As sh1574 almost entirely lacks TRMT2A, treatment with a TRMT2A specific inhibitor should not have any effect on polyQ aggregation. The presence of PolyQ aggregates coincides with cell dysfunction/loss in several model systems [6, 87, 88]. When performing cell viability analysis, sh1574 cells were protected from toxicity induced by expression of HttQ103ex1:GFP compared to shK cells (Figure 6B). Also in the context of polyQ-induced toxicity, a TRMT2A-specific inhibitor should not have any effect on polyQ-induced toxicity in sh1574 cells. Untreated shK cells and sh1574 cells served as controls for inhibitor treatment (Figure 6 A,B).

**Figure 6.**
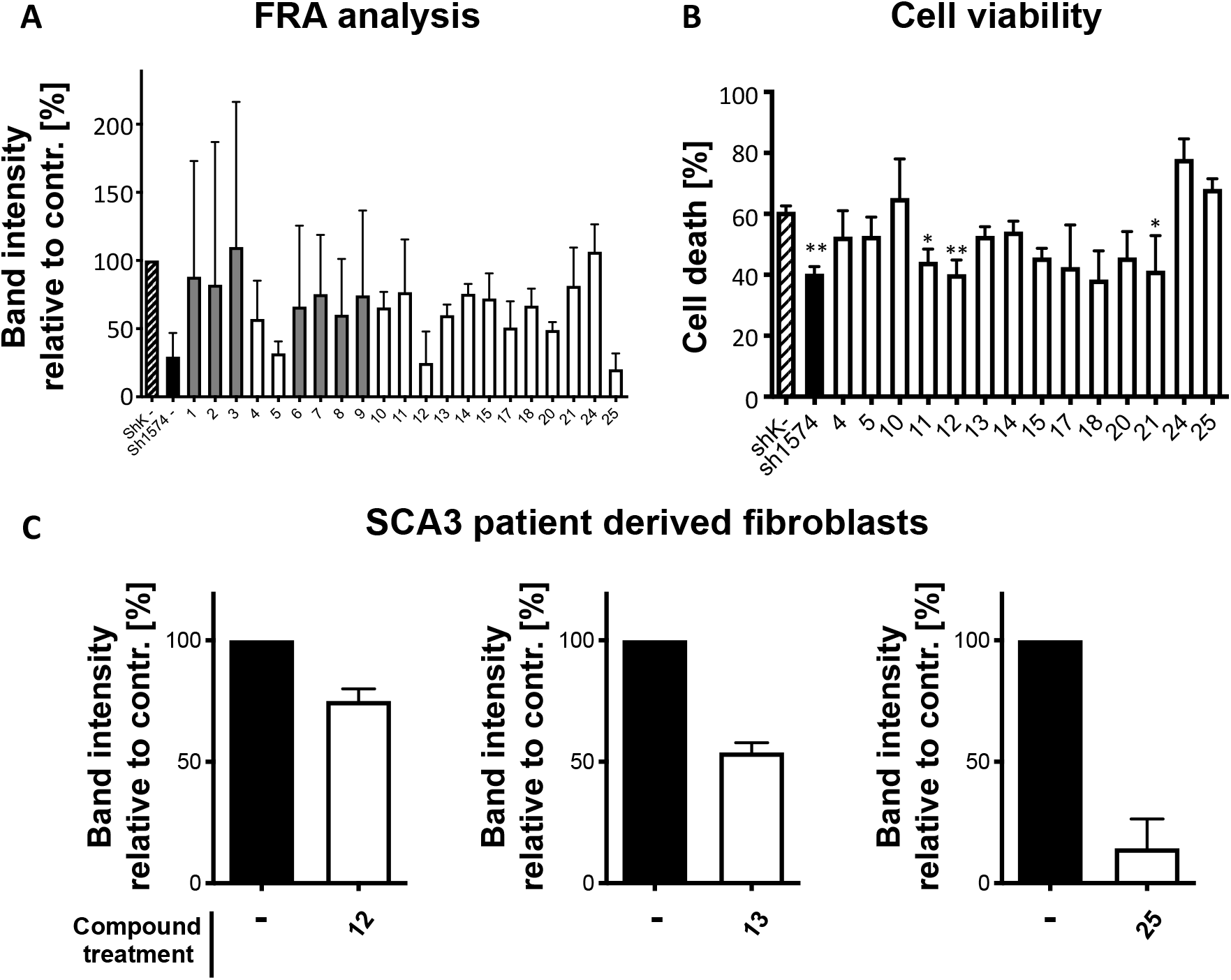
Functional test of potential TRMT2A inhibitors in cells. (A) An FRA analysis was performed to detect the abundance of polyQ aggregates HEK293T cells after inhibitor treatment. The graph depicts a summary of at least three replicated measures. Nonparametric Kruskal-Wallis test followed by Dunn’s multiple comparison test revealed no significant differences. Inconsistent results, as presented by a high SD, are marked (gray). (B) PolyQ-induced cell death was determined in inhibitor-treated HEK293T cells. One-way ANOVA followed by Bonferroni multiple comparison test was used to determine significance. Asterisk indicates significant difference compared to control (shK-); *p<0.05, ** p<0.01. (C) Detection of polyQ aggregates in fibroblasts derived from a HD and a SCA3 patient. Samples were either DMSO treated (contr., black) or treated with indicated inhibitors (white).

ShK and sh1574 cells were transiently transfected with plasmids mediating HttQ103ex1:GFP expression. Four hours post transfection, cells were either treated with inhibitors or with solvent only as control. 48 hours post transfection, brief microscopic inspection was performed to determine the abundance of GFP positive cells. This was done to ensure similar transfection rates among different samples. In addition, Western blot was performed to evaluate the abundance of soluble HttQ103ex1:GFP peptide in transfected shK and sh1574 cells in each experiment. Similar/comparable abundance of soluble HttQ103ex1:GFP peptides in lysates derived from shK and sh1574 cells was a prerequisite to continue with FRA analysis. In FRA we determined whether inhibitor treatment affected aggregate formation in shK cells. As sh1574 cells generally display reduced HttQ103ex1:GFP aggregate load according to a strongly reduced TRMT2A abundance, this served as an internal control. Moreover, sh1574 cells were also treated with respective inhibitors. TRMT2A specific inhibitors were not expected to have any effect on aggregate formation of HttQ103ex1:GFP peptides in sh1574 cells (as these lack the to-be-inhibited protein). Accordingly, the absence of an inhibitory effect of the inhibitor on sh1574 cells implies that the inhibitor specifically exerts its effect via inhibition of TRMT2A and not by the activation of an unrelated mechanism/pathway. Both, a reduction of insoluble HttQ103ex1:GFP in untreated sh1574 cells and the absence of an additional reduction of insoluble HttQ103ex1:GFP in treated cells were a prerequisite for inhibitor selection. An overview of the conducted experiments and outcomes are summarized in Figure S12. In an additional analysis, we tested whether TRMT2A silencing indeed causes reduced levels of m5U. Comparing cell culture supernatants derived from shK cells and sh1574 cells, we detected a significant reduction in m5U abundance in sh1574 by LC-MS/MS (Figure S13).

In this paradigm, 25 compounds were tested in total, of which four were cytotoxic and therefore excluded from further analysis (compounds 16, 19, 22, and 23). The remaining 21 compounds were used for FRA analysis. A summary of FRA analysis for the tested compounds is provided in Figure 6 A. For some compounds (gray), FRA analysis did not provide consistent data, as reflected by the high standard deviation (SD) (SD > 40% of control value). These compounds were 1, 2, 3, 6, 7, 8, and 9. Accordingly, these seven compounds were excluded from further analysis. Among the remaining compounds, three (5, 12, and 25) caused a strong reduction of polyQ aggregate load, comparable to the reduction observed in sh1574 cells (black bar). Seven compounds (4, 10, 13, 14, 17, 18, and 20) caused a moderate to mild reduction of polyQ aggregate load. The remaining four tested putative inhibitors of TRMT2A (11, 15, 21, and 24) did not reduce polyQ aggregation in shK cells (Figure 6 A). Accordingly, 14 compounds (4, 5, 10, 11, 12, 13, 14, 15, 17, 18, 20, 21, 24, and 25) were further analyzed for their effect on polyQ-induced cell death in shK and sh1574 cells (Figure 6 B). Untreated shK cells displayed roughly 60% cell death in our automated cell viability measure based on Alamar Blue staining. In untreated but TRMT2A deficient sh1574 cells, the cell death was decreased by about 20%. Thus, we would expect that shK cells treated with functional TRMT2A inhibitors to display cell death rates of between 40% and 60%. Among the tested compounds, just three (11, 12 and, 18) decreased cell death in a similar way as observed for sh1574 cells. Other compounds even slightly enhanced cell death (10, 24, and 25). The remaining tested compounds gave only mild or no protection towards polyQ-induced cell death.

The overexpression of polyQ peptides fused to GFP has been successfully used to model polyQ aggregation and to induce polyQ toxicity in many model organisms [88–90]. However, in a human disease situation, endogenous polyQ containing proteins are expressed by their native promoter and undergo multiple post-transcriptional modifications like proteolytic cleavage and biotinylation [91]. To further evaluate our compounds as potential therapeutic agents to treat polyQ-diseased patients, we used a patient-derived fibroblast (Figure 6 C). We obtained fibroblasts from patients suffering from SCA3 from the Coriell repository (https://www.coriell.org). Compared to fibroblasts derived from a healthy control, fiborblasts derived from a SCA3 patient displayed no alteration in TRMT2A abundance, as expected. Similarly, spermidine treatment in shK cells did not change the abundance of *Trmt2a* transcripts, indicating that observed treatment effects result from TRMT2A inhibition rather than from alterations in transcript or protein abundance (Figure S14). FRA analysis revealed the formation of RIPA insoluble polyQ aggregates in these cells after two days of culturing. Thus, we applied the same treatment paradigm as for HEK293T cells to test a selection of potential TRMT2A inhibitors in fibroblasts. Compounds were chosen according to their effects in previous analysis and availability. We chose inhibitors 12, 13 and 25. Compounds 12 and 13 reduced polyQ aggregation in FRA analysis and protected HEK293T cells from polyQ-induced cell death. Inhibitor 25 also strongly reduced polyQ aggregation in FRA analysis but it slightly enhanced polyQ-induced cell death. Treatment with these inhibitors decreased the abundance of polyQ aggregates in SCA3-patient-derived fibroblasts (Figure 6 C).

Compounds 4, 9, 10, 11, and 12 were previously reported in the literature as inhibitors of *E. coli* or *Saccharomyces cerevisiae* homologs of TRMT2A, respectively [92, 93]. Neither L-norleucine (compound 4), L-methionine (compound 9), nor L-ethionine (compound 10) was able to reduce cell death, suggesting a considerably different catalytic binding site. The two polyamines tested, putrescine (compound 11) and spermidine (compound 12), but also the organomercury agent p-chloromercuribenzoic acid (compound 5, PCMB), which reacts with free thiol groups in proteins containing cysteine, lowered aggregate abundance and cell death. Interestingly, the RRM of TRMT2A features a solvent-exposed cysteine (C111). Polyamines are known to bind tRNA methyltransferases and, depending on Mg^2+^ concentration they can either behave as activator [92] or inhibitors [93]. In our experimental set-up, they appear to act as inhibitors.

Of note, compounds 13-25 resulted from structure-based drug discovery and the putative cryptic allosteric pocket on the helical site of the RRM. The majority of these compounds were predicted to bury at least one aromatic moiety into the cryptic pocket. Compounds 16, 19, 22, and 23 are cytotoxic. Compound 19 was used as an internal negative control, as it contains a bulky and hydrophobic naphthalenyl group, which does not fit into the cryptic site based on our docking study. Not only does compound 19 fail to rescue the cell lines affected by polyglutamine toxicity but it also reveals additional cytotoxicity.

Intriguingly, ligands featuring a phenyl group (e.g., compounds 13-15, 17, 18, 21, and 25) show an effect in at least one assay. We speculated that if these compounds bind indeed into the cryptic site, their phenyl group might take the place of the indole ring of W134 (as seen in Figure 1B), and then could hereby form π-stacking interactions not only with F92 but also with the indole ring of W134, now slightly displaced with respect to the X-ray structure (see Figure 4 B).

Furthermore, since we did not measure TRMT2A inhibition directly, but by means of an assay, we wanted to investigate the possibility of pan-assay interference compounds (PAINS filter, see methods). Compounds, 18, 19, 20, and 21, did not pass the PAINS filter. As previously mentioned, compound 19 increased cytotoxicity. Nevertheless, the remaining three compounds (18, 20, and 21), might be false positive hits.

We also evaluated the possibility of off-targets binding with a reversed *in silico* screening approach, as implemented in SwissTargetPrediction, [94] to elucidate putative human protein targets for our non-polyamine bioactives. For these compounds, the most frequently predicted target class was kinases. However, this result may be biased due to the great number of known kinase inhibitors.

Notably, some of the compounds discovered by structure-based discovery (13, 15, and 17) along with the polyamines (spermidine and putrescine), reported in the literature as inhibitors for TRMT2A homologs, consistently and effectively lowered cell death and aggregation in polyQ affected HEK cells and do not contain critical substructures that often confound certain cell assays. Finally, compounds 13 and 21, which show structural similarity in the form of a phenyl group, drastically reduced aggregate formation in SCA3 patient derived fibroblasts (see Figure 6 C) with improved predicted solubility while retaining the predicted blood-brain-barrier permeability [95].

### 3.5 Biophysical measurements of TRMT2A RRM and compounds

The compounds tested on SCA3 patient fibroblasts (12, 13, and 25) and the two additional compounds that showed promising results in cell death and FRA analysis (15 and 17) were selected for further *in vitro* interaction experiments. To analyze RRM-compound interaction, surface plasmon resonance (SPR) experiments were performed. The polyamine spermine was used as a positive control. Polyamines are known to bind tRNA methyltransferases [92, 93]. Indeed, we observed that spermine binds in the micromolar range to the TRMT2A RRM (K_D_ = 1.30 +/− 0.27 μM; Figure 7). In contrast, none of the selected compounds showed any binding to the protein (Figure S15), with the exception of compound 12, the polyamine spermidine (K_D_ = 2.28 +/− 0.69 μM; Figure 7). Noteworthy, SPR experiments were performed with low salt concentrations in the running buffer to also allow for the occurrence and detection of lower affinity binding events. The size difference between compounds (> 0.2 kDa) and the TRMT2A RRM (9.2 kDa) was taken into consideration by coupling high amounts of protein (3000 RU) to the CM5 chip. Taken together, except for polyamines spermine and compound 12/spermidine, no direct binding to the RRM could be detected.

**Figure 7.**
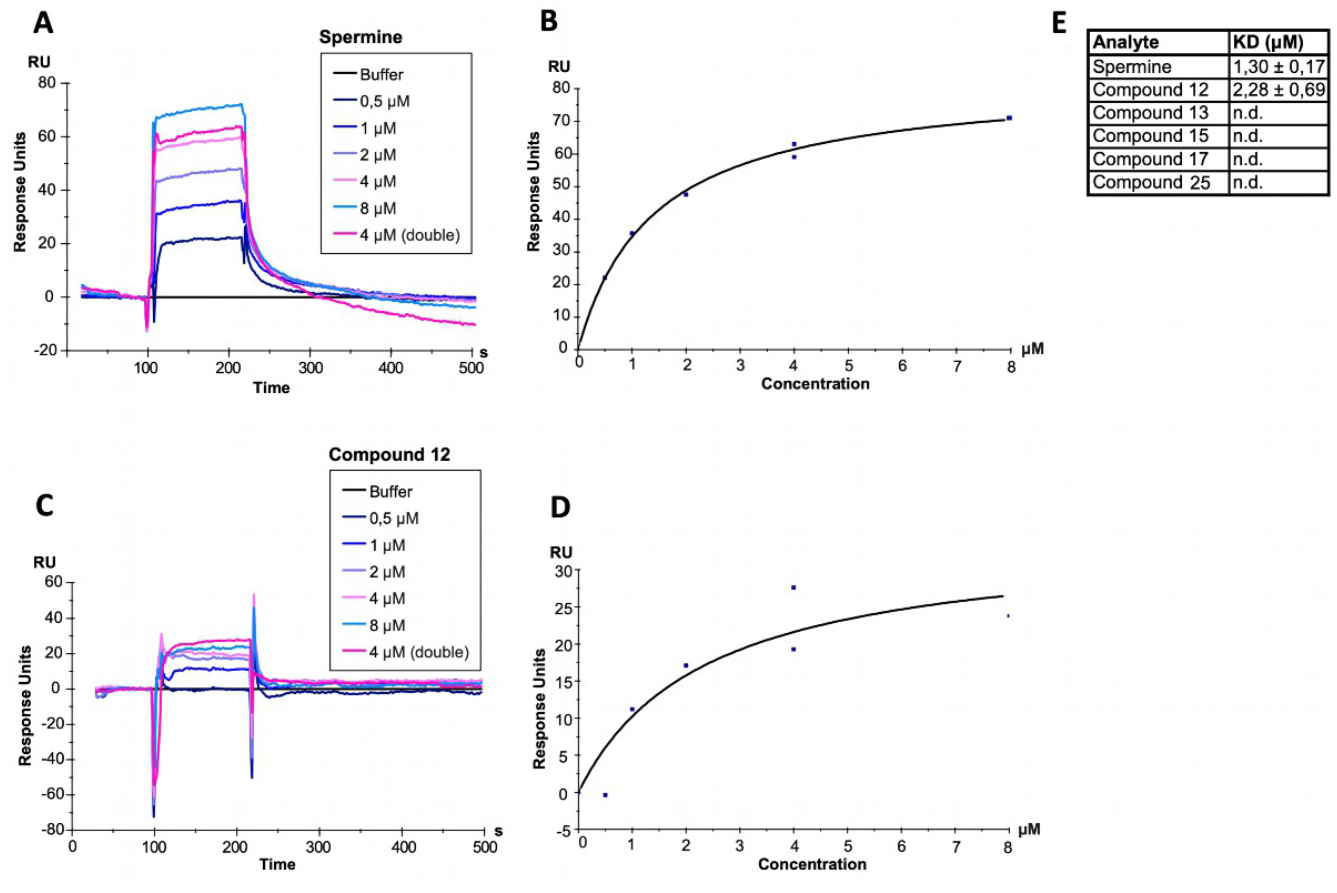
Results of Surface Plasmon Resonance experiments. (A-C), The TRMT2A RRM was coupled to a CM5 chip reaching 3000 RU. A concentration series from 0.5 to 8 μM spermine (A)/compound 12 (C) was injected, resulting in the illustrated, double-referenced binding curves. A representative curve from a triplicate measurement is shown. (B-D),The response at the equilibrium of the binding curve is plotted against spermine (B)/compound 12 (C) concentration. (E) A steady-state affinity model was fitted and allowed for the calculation of K_D_ values.

## 4. Conclusion

In this work, several *in silico* strategies were performed to identify a small library of ligands able to modulate the activity of tRNA methyltransferase 2 homolog A (TRMT2A) in cells.

TRMT2A is an enzyme that converts uridine to m^5^U_54_ on tRNA across species, from yeast to man [15]. Unfortunately, very little is known concerning the role of this modification in mammals [13]. It was shown that an absence of TRMT2A reduces polyQ aggregation and polyQ-induced toxicity in HEK293T cells [12]. This suggests that reducing the abundance of the tRNA modification m^5^U_54_ might have functional consequences in the context of polyQ-induced toxicity. Thus, it was reasonable to test known and new (putative) inhibitors of TRMT2A with regard to their effect on polyQ aggregation and polyQ-induced toxicity. In total, we assayed 25 potential inhibitors. While four compounds turned out to be cytotoxic, the ten other compounds reduced polyQ-aggregation in FRA. Amongst the latter, three compounds (5, 12, and 25) turned out to reduce polyQ-aggregation very efficiently. The observed effect on polyQ aggregation was as strong as after *TRMT2A* silencing.

Due to its similarity to yeast Trm2p, mammalian TRMT2A harbors a putative RRM in the N-terminus and the CD at the C-terminus [13]. Inhibiting its function by directly interfering with the CD activity or with TRMT2A binding to the RNA might ameliorate some pathological effects linked to polyQ toxicity. No full-length structure is available for this protein. Here we managed to crystallize and solve the structure of a sub fragment of TRMT2A corresponding to its RRM. We conducted two parallel strategies for finding inhibitors; one exploiting the structural information of the crystal structure of the TRMT2A RRM, the other exploiting the available knowledge of general methyltransferase inhibitors. The RRM is characterized by the absence of protein pockets. However, we identified a putative cryptic site, potentially able to allosterically affect RNA binding. As a first strategy, we conducted a structure-based virtual screening, and the selected candidates were tested both *in vitro* and *in cell* experiments. Several compounds with similar substructures induced a reduction of polyQ aggregation *in cell*. Also, two of them, 13 and 25, were able to decrease polyQ aggregates in SCA3-patient-derived fibroblasts.

For the CD, a pharmacophore model was identified and used for a ligand-based virtual screening. Also, in this case, several selected candidates were able to reduce PolyQ aggregation in cell culture. Moreover, the polyamine spermidine (compound 12) caused a decrease in the number of polyQ aggregates in SCA3-patient-derived fibroblasts and directly interacted with the RRM in SPR experiments. However, spermidine is predicted to have an unspecific binding in the RRM, namely to cluster around negatively charged and solvent-exposed side-chains, as similar in-size and in-volume fragments does (Figure S16).

Except for spermidine, we were not able to prove any direct binding between ligands screened against the TRMT2A RRM and this domain. There are several possible explanations for this. Ligands targeting the RRM were designed for the cryptic site in the *minor/small groove*. The lack of observed interaction of the compounds with the RRM could be due to the possibility that the cryptic site might require the presence of tRNAs for its formation. Furthermore, the rest of the protein might also play an important role in the suggested allosteric mechanism, connecting the tRNA binding site with the identified cryptic site. In addition, some studies show that not all of the cryptic pockets that are seen to open during MD simulations are capable of binding a ligand with substantial affinity [96]. Our model of allosteric communication between the tRNA binding site and the cryptic pocket relies on a normal mode analysis. This approach involves drastic approximations and only provides reasonable prediction if correlated thermal motions of these two areas exist [97]. Also, ligands could target the CD or other protein regions that were excluded in our assays. However, we consider this option the least likely as the *in-silico* selection of ligands was based on features derived from the experimental crystal structures of the TRMT2A RRM.

Another possible explanation is that the observed effects of inhibitors in cells might be the result of an interaction with other protein/s and/or activation of cellular pathways compensating aggregation-induced detrimental effects. In an attempt to exclude this possibility, we tested our ligands also on TRMT2A-silenced cells (sh1574), and no effect on aggregation or cell death was observed, pointing toward a TRMT2A-dependent effect of our ligands.

As mentioned above, spermidine was shown to bind the TRMT2A RRM. Like other polyamines, this compound has a well-proven affinity for nucleic acids as well as proteins. Since polyamines have been implicated in translational control and RNA binding [98], it is not entirely surprising to see also binding to the RRM. It remains to be shown how specific the interaction between the TRMT2A RRM and spermidine really is.

No therapy has been approved to interfere with the clinical progression of these diseases in humans [9]. Moreover, ASOs, a pivotal therapeutic venue that raised hope in the HD community, have unfortunately failed in a late-stage clinical trial [11]. Moreover, ASOs development has previously been hampered by invasive administration requirements and long optimization strategies [99]. Therefore, targeting a novel CAG-disease modifier such as TRMT2A may still represent an alternative therapeutic approach with several advantages over ASO-based therapies..

## Acknowledgment

We like to thank Vera Roman for her technical support. This work was supported by a grant by the Federal Ministry of Education and Research (BMBF) to AV and DN (The PolyQure project: grant numbers 16GW0306K, 16GW0307K, respectively). This research was supported by the Joint Lab “Supercomputing and Modeling for the Human Brain”. Structure determination was done with support of the X-ray crystallography platform at the Institute of Structural Biology, Helmholtz Zentrum München.

## Supplemental Material

Figures S1–S16, Table S1-S2, and Text Supplements S1-S4

## Notes

### Competing Interest Statement

The authors have declared no competing interest.

