## Supplemental Material for "Small-molecule modulators of TRMT2A decrease PolyQ aggregation and PolyQ-induced cell death"

† = shared first author

*= shared senior authorship

### SUPPLEMENTAL FIGURES

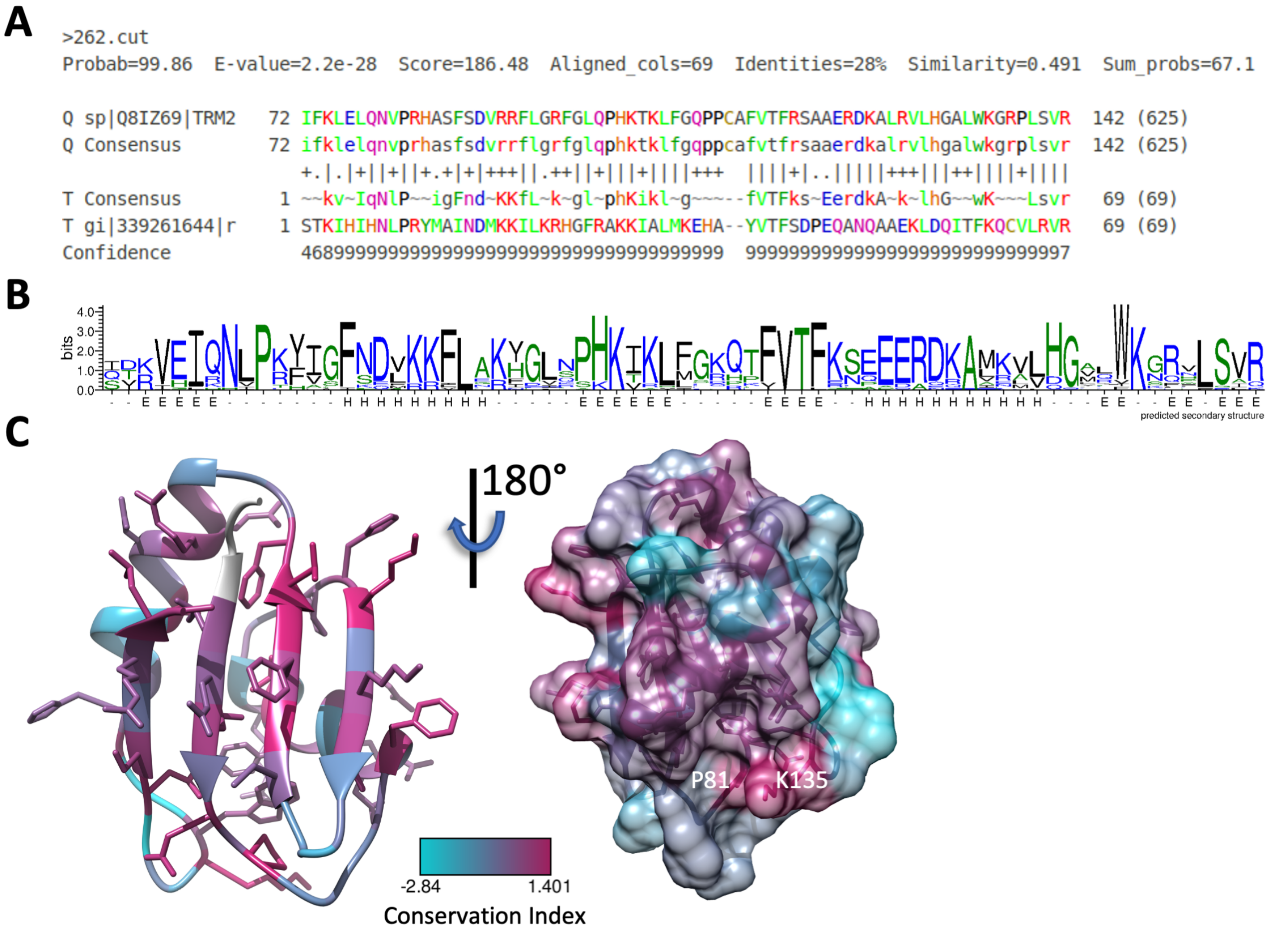

**Figure S1. Conservation.** (A) Multiple Sequence Alignment (MSA) of several RRMs related to TRMT2A (Q8IZ69, retrieved from the RRMdb[1]); These include *Trichinella spiralis, Caenorhabditis elegans, Caenorhabditis brenneri, Condylura cristata, Dasypus novemcinctus, Callorhinchus milii, Ornithorhynchus anatinus, Xenopus (Silurana) tropicalis, Gallus gallus, Chaetura pelagica, Latimeria chalumnae, Anas platyrhynchos, Alligator mississippiensis, Cariama cristata, Tauraco erythrolophus, Anolis carolinensis, Ophiophagus hannah, Monodelphis domestica, Sarcophilus harrisii, Lepisosteus oculatus, Danio rerio, Cynoglossus semilaevis, Esox lucius, Neolamprologus brichardi,* and *Oryzias latipes*. (B), Weblogo[2] depiction of the MSA. The overall height of the stack indicates the sequence conservation at that position, while the height of each amino acid within the stack indicates the relative frequency of each amino at that position. (C), Entropy-based conservation indices[3] mapped onto the crystal structure of the TRMT2A RRM (left: protein view with the RNA binding site on the top, right: protein view obtained by rotating the protein's orientation shown on the left by 180 degree). Higher indices reveal a higher degree of conservation, which is not only found on the putative tRNA binding site (in front of the β-sheet, left) but also on the helical back (conservation index mapped onto the Van der Waals surface, right), and in particular on P81 and K135. Created with UCSF Chimera [2].

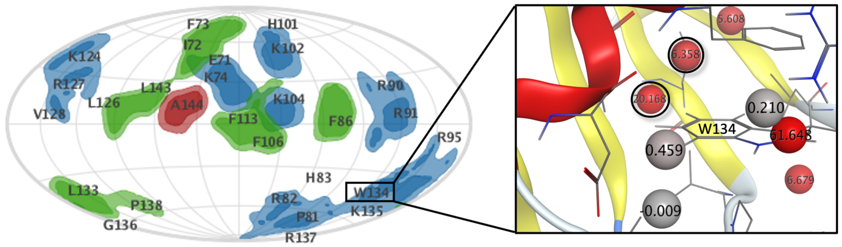

**Figure S2. Water Analysis.** Protein patches projected onto a sphere encapsulating the 2 Å resolution structure of the TRMT2A RRM are outlined as blue areas (positive), red (negative), and hydrophobic sites (green) with key residues labeled. On the right, a zoom-in around W134 with solvent analysis using 3D-RISM is shown. Spheres represent predicted hydration site centers and are labeled according to computed free energies. Red sites indicate propensity peaks of unstable water, and gray sites indicate stable ones. Two unstable water propensity centers are buried 2.5 Å beneath W134 (highlighted with black circles). Depicted with MOE (Molecular Operating Environment) Version 2019.01. Chemical Computing Group, (CCG).

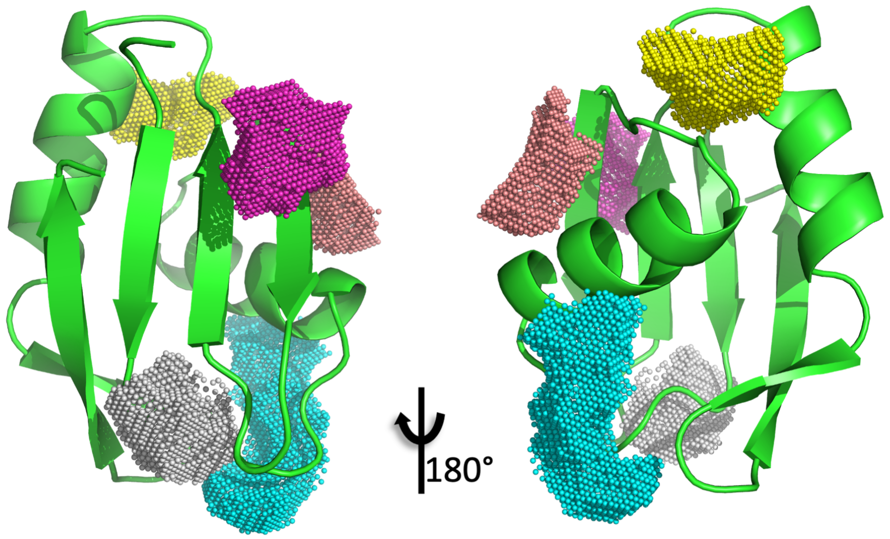

**DoGSite Pocket Assessment**

| *Name* | *DrugScore* | *SimpleScore* | *Volume (Å³)* |
|---|---|---|---|
| **P1 (cyan)** | 0.18 | 0.16 | 221.44 |
| **P2 (magenta)** | 0.09 | 0.22 | 153.66 |
| **P3 (yellow)** | 0.44 | 0.21 | 104.19 |
| **P4 (salmon)** | 0.26 | 0 | 100.86 |
| **P5 (white)** | 0.36 | 0.08 | 100.86 |

**Figure S3.** Binding sites predicted with DoGSiteScorer[4] based on the protein crystal structure of the RRM of TRMT2A with 2.0 Å resolution. None of the outlined sites form a deep cavity. Since *in silico* predictions of small molecule ligands for such sites would likely yield only modest binding affinities, we did not model ligands for these sites.

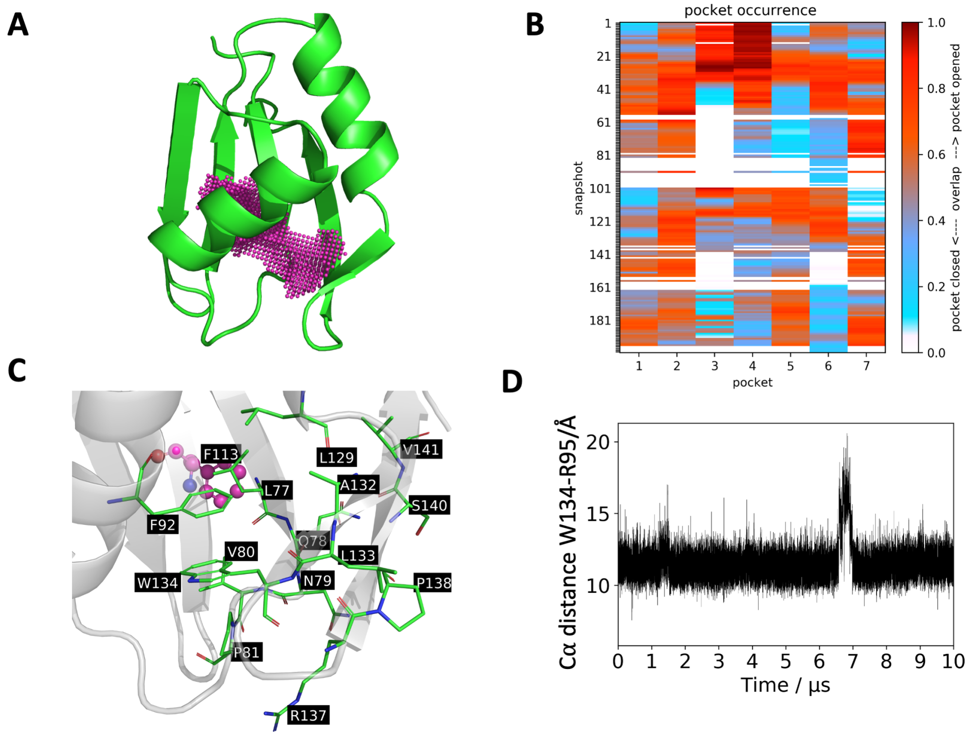

**Figure S4. Structural analysis of the high-resolution crystal structure of the TRMT2A RRM.** (A), DoGSiteScorer[4] prediction of a binding site at the helical back of the high-resolution RRM structure. This cavity not only overlaps with the small groove in the 2.0 Å structure but also protrudes deep into the domain, which is consistent with our previous observation that there might be a cryptic pocket in this region. However, a DrugScore [5] < 0.01 and volume of 153.1 Å$^3$ indicated this pocket as equally undruggable. (B), K-means clustering of the geometric centers of the side chain atoms with TRAPP [6] at a pocket occurrence of 25 % yields seven subpockets in the *minor groove* region. In accordance with the 2 Å resolution structure results, subpockets are opening and collapsing during the implicit solvent MD. (C), The AlloPred[7] analysis also indicated allosteric communication between F113 (magenta ball-sticks rep.) and residues (e.g., (green sticks, W134) in the *minor groove* region. (D), For the 2 Å resolution crystal structure, the centroids distances of R95 and W134 indicate unfolding/refolding, starting after approx. 6.5 μs.

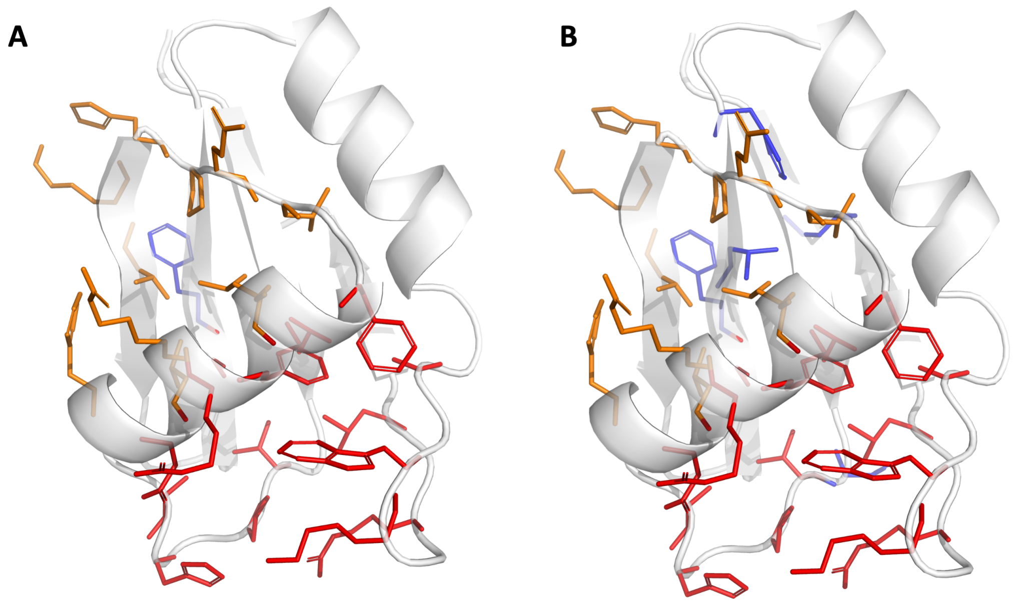

**Figure S5. AlloPred analysis of the 2 Å crystal structure of the TRMT2A RRM.** The crystal structure of the RRM is represented as a white transparent cartoon. Residues depicted as blue sticks were used for the orthosteric site (e.g., tRNA binding site) definition. Red and orange sites were detected as possible allosteric sites. (A), Only the solvent-exposed F113 on the β-sheet for the definition of the orthosteric site was used. (B), All conserved residues on the β-sheet were used to define the orthosteric site in AlloPred[7]. Two allosteric sites were detected (red, orange sticks) independently of the orthosteric-site definition. Interestingly, the predicted allosteric effect increases when the tRNA interaction site definition is extended from the canonical RNP1 phenylalanine to all conserved RNP1/2 aa. The same eight allosterically communicating aa are found, whereas the inter-pocket distance decreases to 11.7 Å. The solvent-exposed hydrophobic F113 site chain is known to form π-π-stacking interactions with tRNA nucleotides[8].

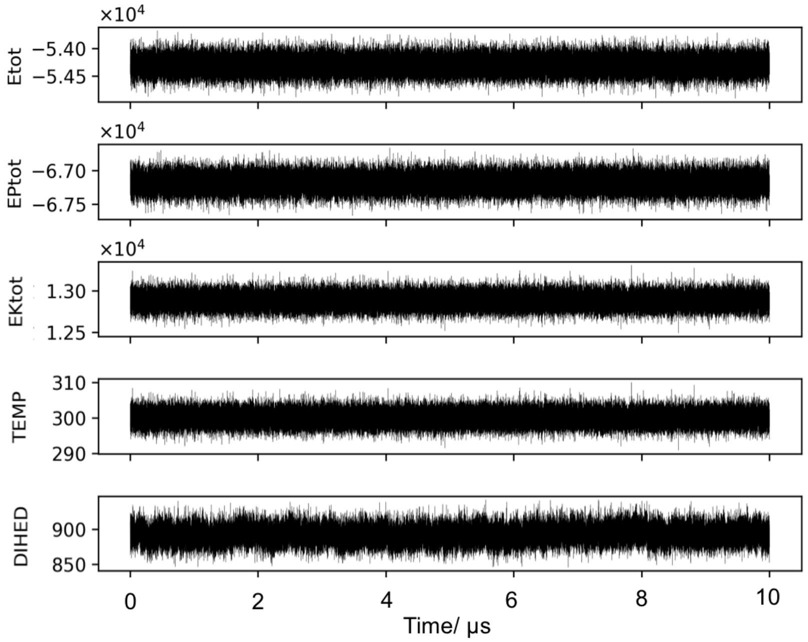

**Figure S6. Simulation quality assessment.** the fluctuations of several simulation parameters are shown. From the top: Etot (total energy), EPtot (potential energy), Ektot (kinetic energy), TEMP (Temperature in K), and DIHED (Dihedral energy). Energies in kcal/mol. Temperature fluctuations are within 10 K and all other parameters suggest a stable simulation.

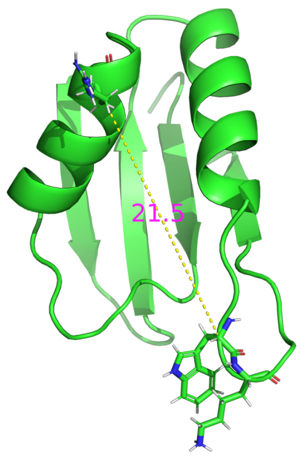

**Figure S7. Selected MD snapshot.** MD snapshot after 2.7776 μs of simulation time: We observed a $C_\alpha$ distance between R95 (top) and W134 (bottom) of 21.5 Å (yellow dashed line). This conformation seems to be stabilized by a cation-$\pi$ interaction between W134 and K135.

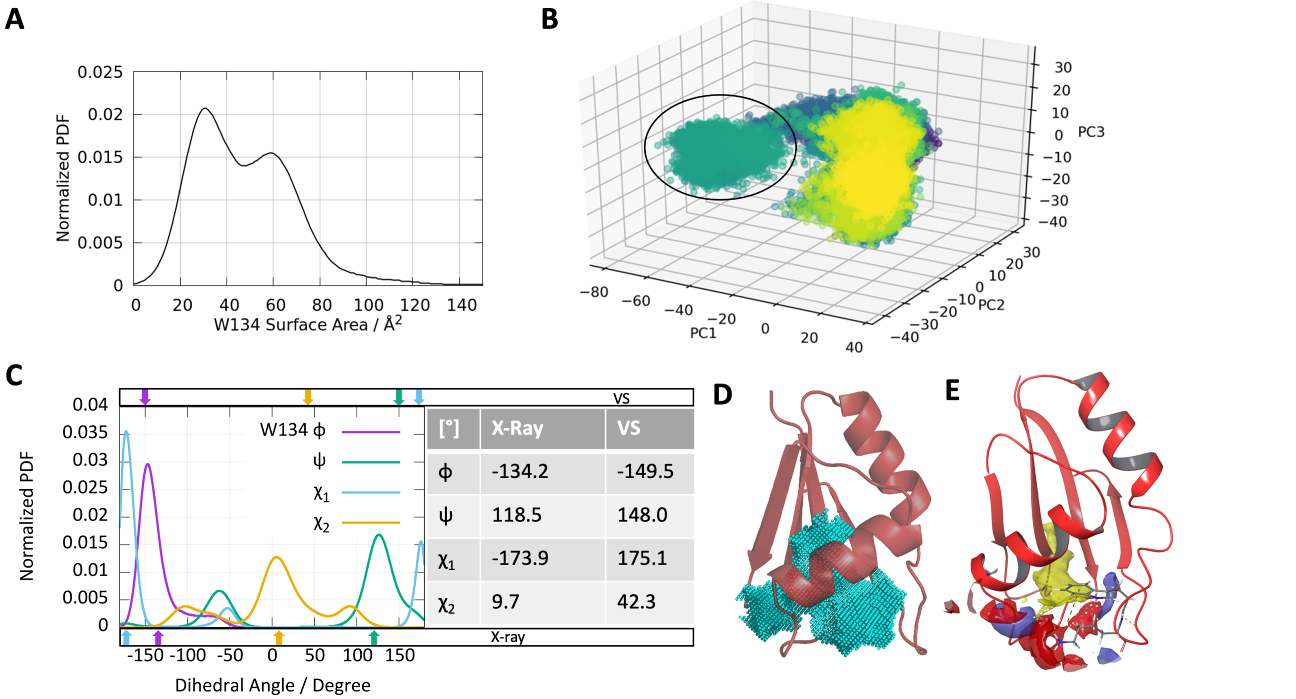

| [°] | X-Ray | VS |
|---|---|---|
| ϕ | -134.2 | -149.5 |
| ψ | 118.5 | 148.0 |
| $\chi_1$ | -173.9 | 175.1 |
| $\chi_2$ | 9.7 | 42.3 |

**Figure S8. W134 and the cryptic pocket.** (A), Structural changes in the RRM are further highlighted by the normalized solvent accessible surface area histogram of W134 with a Gaussian kernel density estimator (KDE). W134 is mostly buried, consistent with its overall hydrophobic side chain. However, since the distribution is roughly bimodal W134 seems to be encountered also frequently in a more exposed state. (B), Principal component analysis (pytraj version 2.0.2) distinguishes the conformations between ~5 to 6 µs, corresponding to the highlighted fraction of frames. (C), Comparison between the X-ray structure (low resolution) and the frame selected for the VS using dihedral angle histograms for W134 and overall propensities using KDE during the simulation. Here, $\chi_1$ changes from the X-ray to the MD frame for virtual screening (VS) from -173.9° to 175.1. (D), DoGSiteScorer[4] results of the frame selected for VS and the investigated pocket. (E), Physiochemical analysis with Schrödinger SiteMap of the frame used for VS (SiteScore 1.13 and DScore 1.16). SiteMap[9] grid points are white, hydrophobic maps in yellow, hydrogen bond donor and acceptor maps in blue and red, respectively.

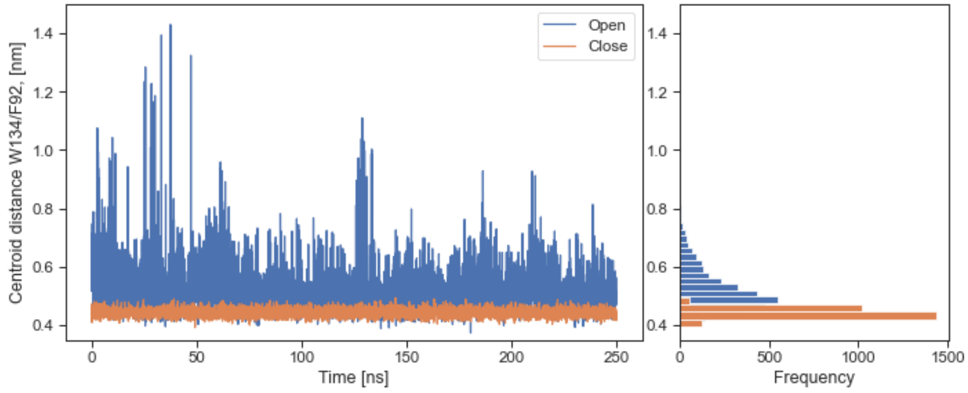

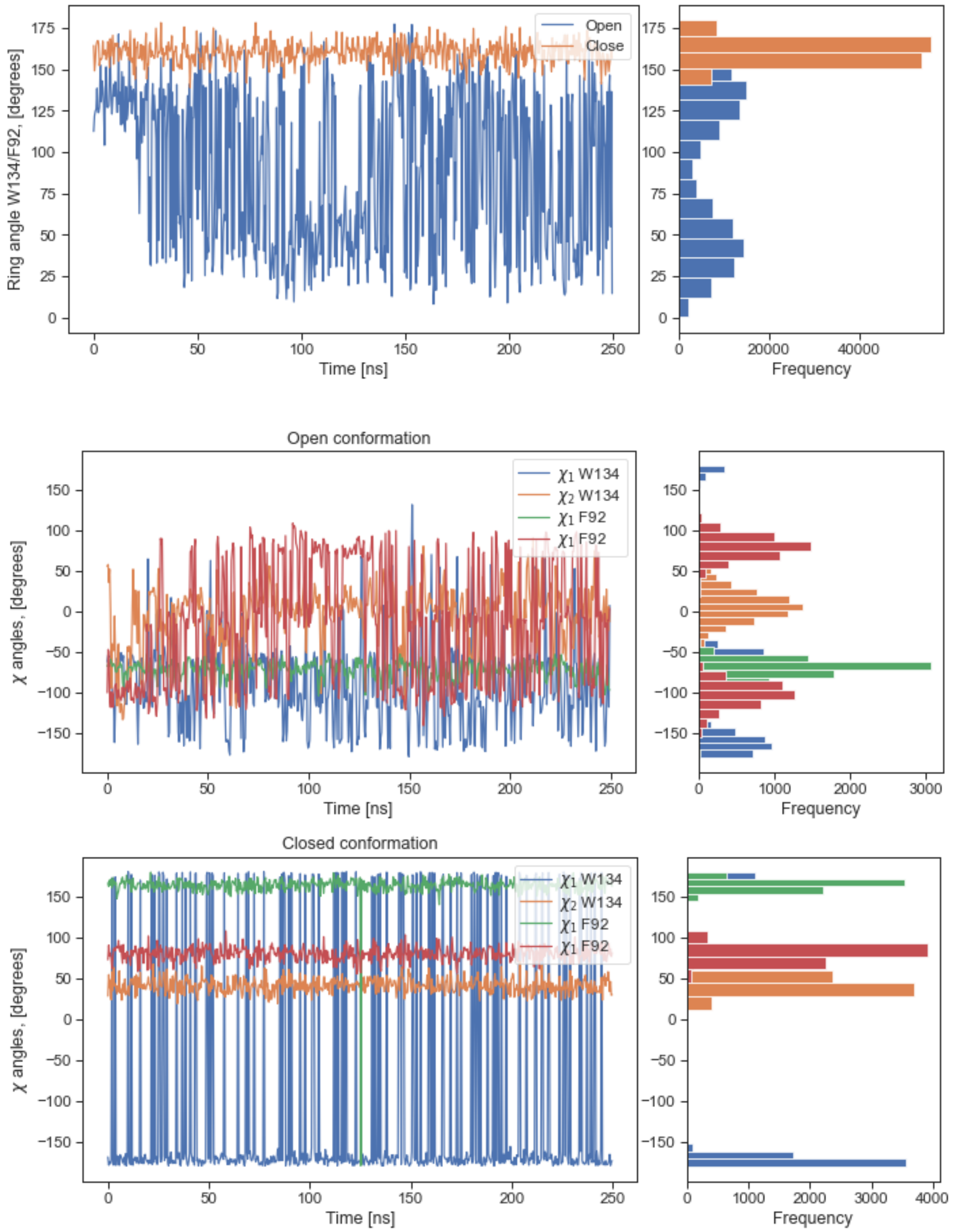

**Figure S9. REST2 simulations.** Ring centroid distances and angles between the F92 and W134 side chains from the REST2 simulations as function of time. The overall frequency of each quantity is also reported for each plot.

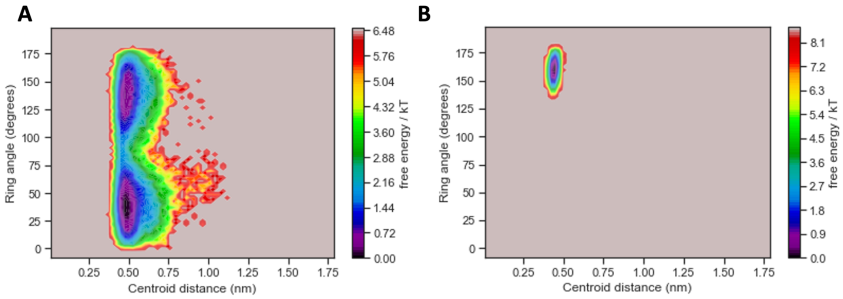

**Figure S10. Conformational landscape.** Conformational landscape using ring centroid distances and angles between the F92 and W134 side chains starting from the open (A) and close (B) conformations.

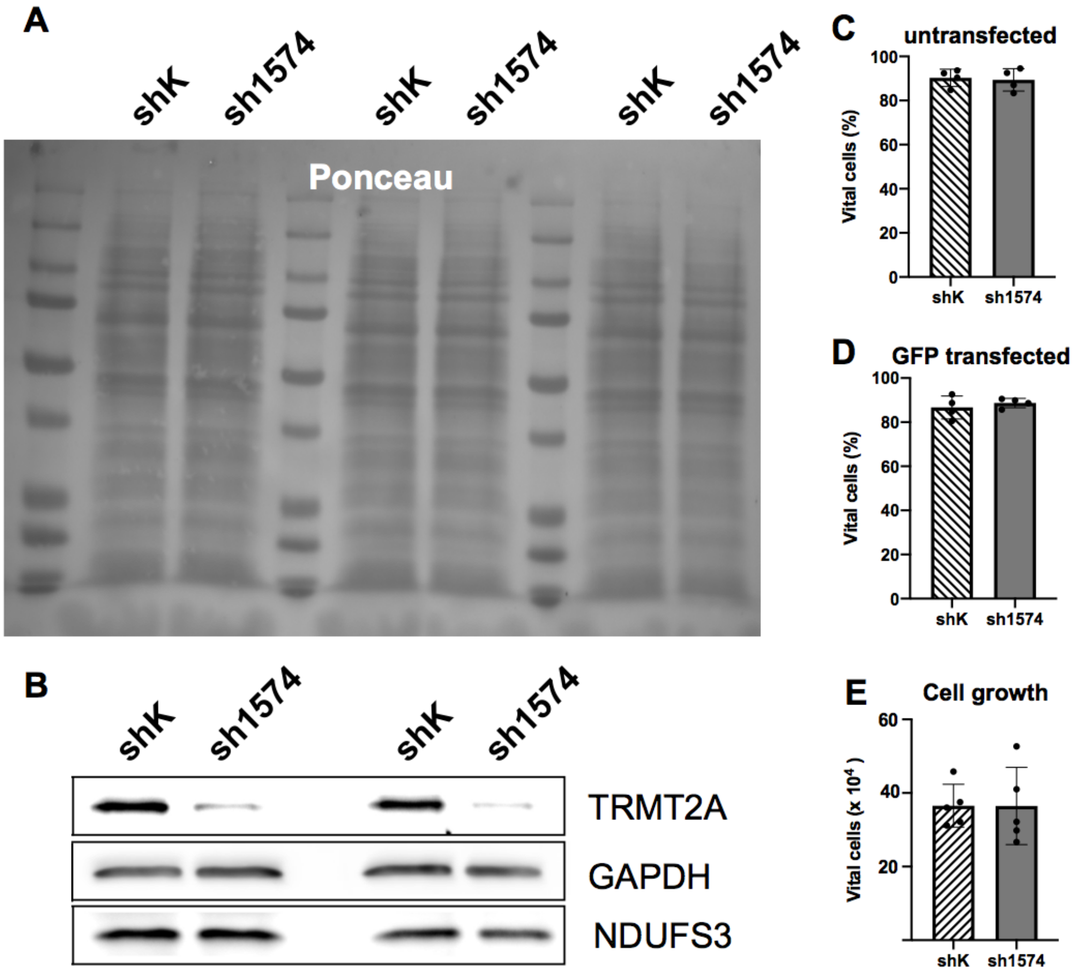

**Figure S11: Analysis of TRMT2A deficient (sh1754) and control cells (shK) with regard to overall protein abundance and cell viability**

Equal amounts of cell lysates (normalized to cell number, 1 million cells) derived from control (shK) and TRMT2A-silenced (sh1574) were submitted to SDS-PAGE and blotted onto a nitrocellulose membrane. A, Ponceau staining of the membrane revealed no obvious difference in the abundance of overall protein. B, Western blot analysis was performed with specific antibodies to determine the abundance of TRMT2A, Glycerinaldehyd-3-phosphat-Dehydrogenase (GAPDH) and iron-sulfur protein components of mitochondrial NADH:ubiquinone oxidoreductase (NDUFS3). Bar graphs depicting relative amount of vital cells (as per cent of total cells) in (C), untransfected and (D), GFP-transfected shK cells and sh1574 cells 48 h post seeding. E, Bar graph depicts amount of shK cells and sh1574 cells (seeded at a density of 250000 per well in a 6-well plate; 28000 cells/cm2) counted 24 h post seeding. No significant difference (C-D; non-parametric Mann-Whitney test (Two-tailed) p>0.05) was detected comparing shK and sh1574 cells.

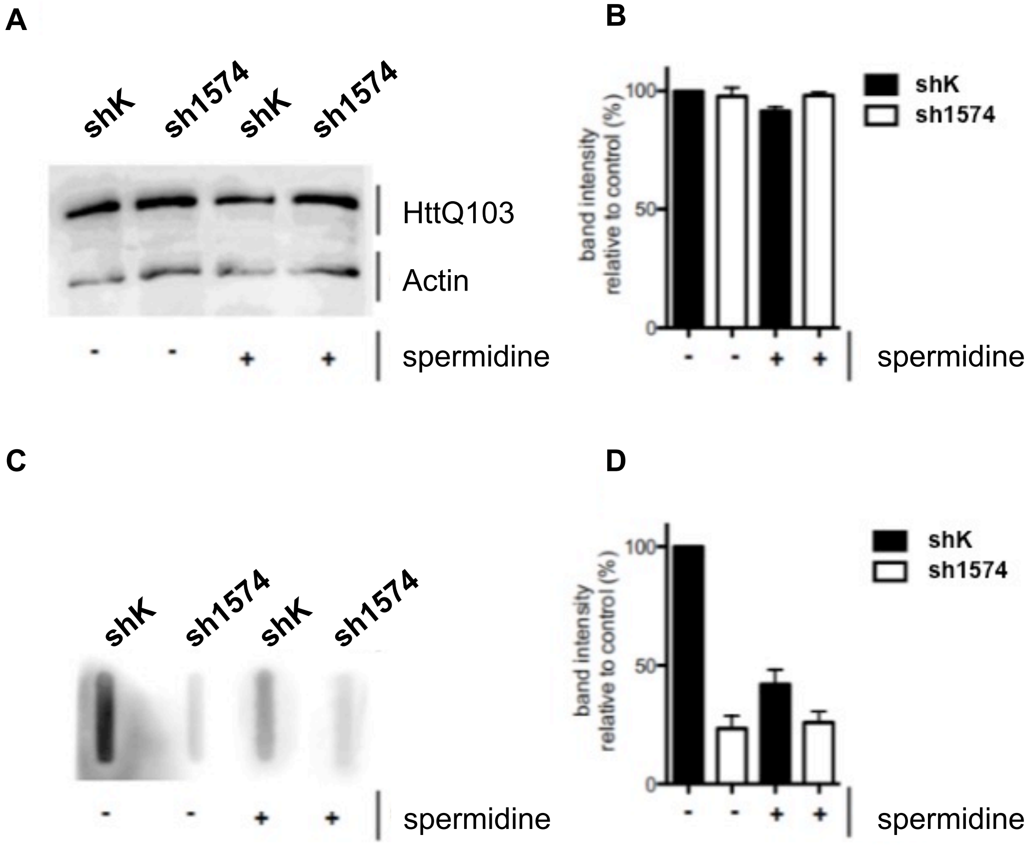

**Figure S12: Analysis of the amount of aggregates of GFP-HttQ103 in spermidine treated TRMT2A deficient (Sh1754) and control cells (ShK)**

(A), Western blot analysis of cell lysates, derived from control (ShK) and TRMT2A-silenced (sh1574) cells, transfected with HttQ103:GFP with or without spermidine treatment. The abundance of HttQ103:GFP was analyzed using an antibody specific to GFP. Determination of Actin abundance was used to show equal protein load in both conditions. (B), Bar graph displays quantification of WB-band intensities derived from three biological replicates. No differences in HttQ103:GFP abundance were observed. (C), Filter retardation assay was performed to determine the abundance of polyQ aggregates. A specific antibody against elongated PolyQ was used for detection. (D), Bar graph representing polyQ band intensities derived from three biological replicates.

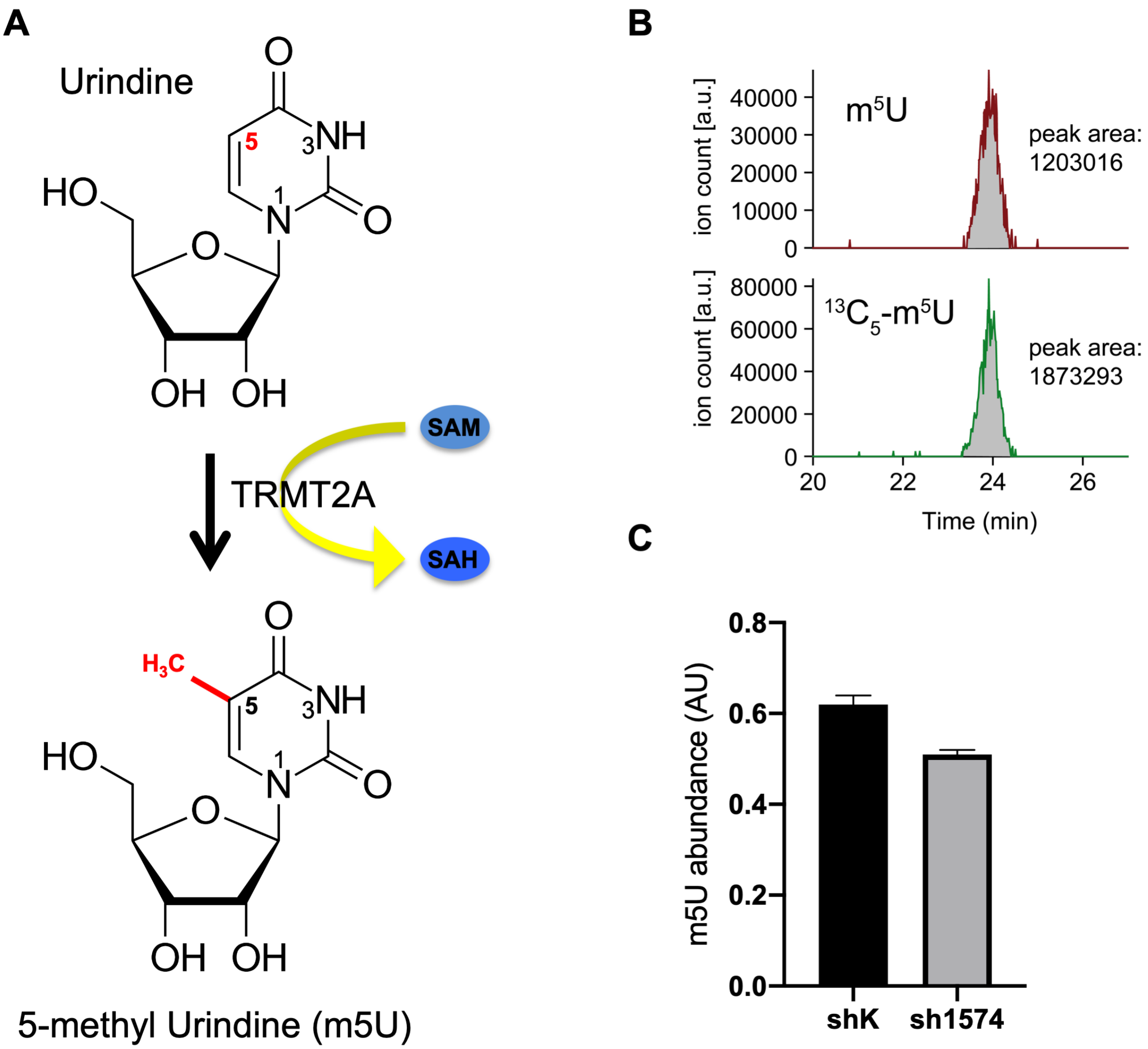

**Figure S13**. **m5U generation and detection**. (A), Cartoon visualizing the generation of Uridine to 5-methyl-Uridine (m5U) by TRMT2A enzymatic reaction. TRMT2A is a S-adenosylmethionine (SAM) dependent methylase that uses SAM as a methyl donor for the methylation of Uridine. Products of this reaction are 5-methyl Uridine (m5U) and S-adenosylhomocysteine (SAH). (B), Extracted ion chromatograms of natural m5U (top) and its 13C5-labeled isotopologue (bottom) after LC-ESI-MS analysis of a sample. Peak area ratios were used to determine the relative m5U-content of the samples. (C), Determination of m5U abundance in cell culture supernatants derived from shK and sh1574 cells using LC/MS. Ordinary unpaired t-test (two tailed) was used determine significance. p* < 0.05.

**A**

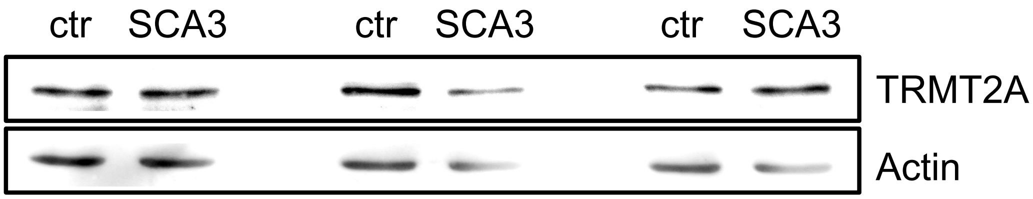

**B**

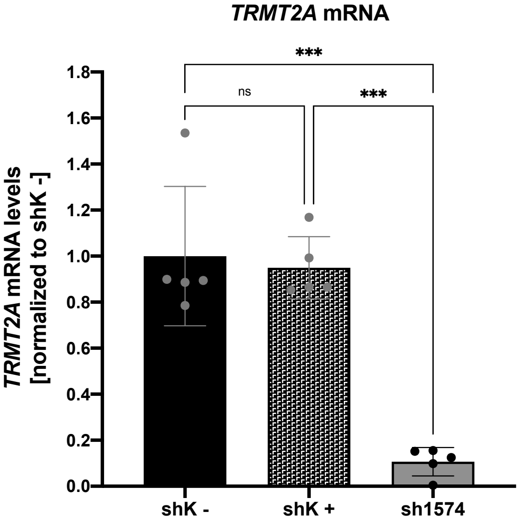

**Figure S14: TRMT2A abundance comparison in fibroblasts of SCA3 patients and healthy control as well as in treated and untreated shK cells**

(A), Western blot analysis detecting TRMT2A abundance in fibroblast lysates derived from a healthy control (ctr) or a SCA3 patient. Detection of Actin was used for normalization. (B), qRT-PCR data partially depicted in Fig. 5 D. Bar graph summarizes qRT-PCR analysis of TRMT2A transcript levels in solvent treated shK cells (-) and spermidine treated shK cells (+) as well as in untreated sh1574 cells. One-way-ANOVA followed by post-hoc Tukey's multiple comparisons test was used to determine significance; ***$p<0.001$, ns = not significant.

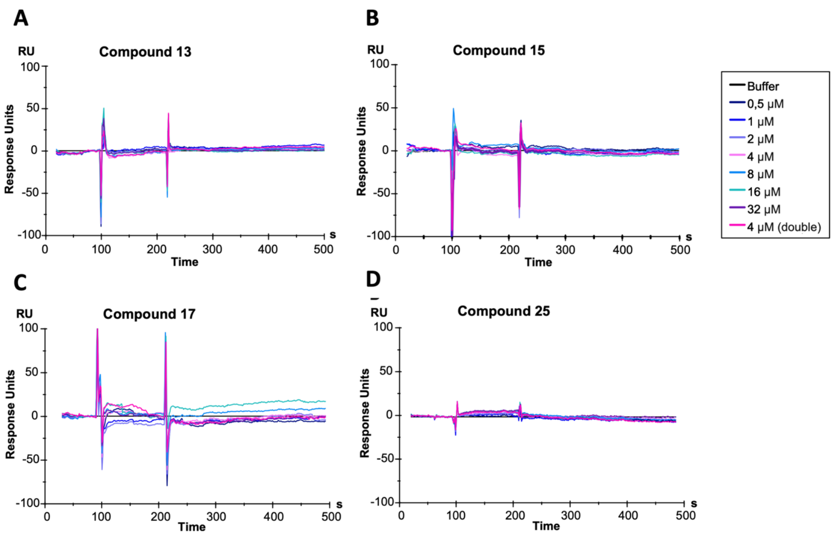

**Figure S15. SPR experiments.** Surface Plasmon Resonance results for compounds 13, 15, 17, and 25. For all experiments, the TRMT2A RRM was coupled to a CM5 chip reaching 3000 RU. A concentration series from 0.5 to 32 µM compound solution was injected, resulting in the double-referenced curves illustrated. (A), compound 13, (B), compound 15, (C), compound 17, and (D), compound 25 do not show any binding. Representative curves from triplicate measurements are shown.

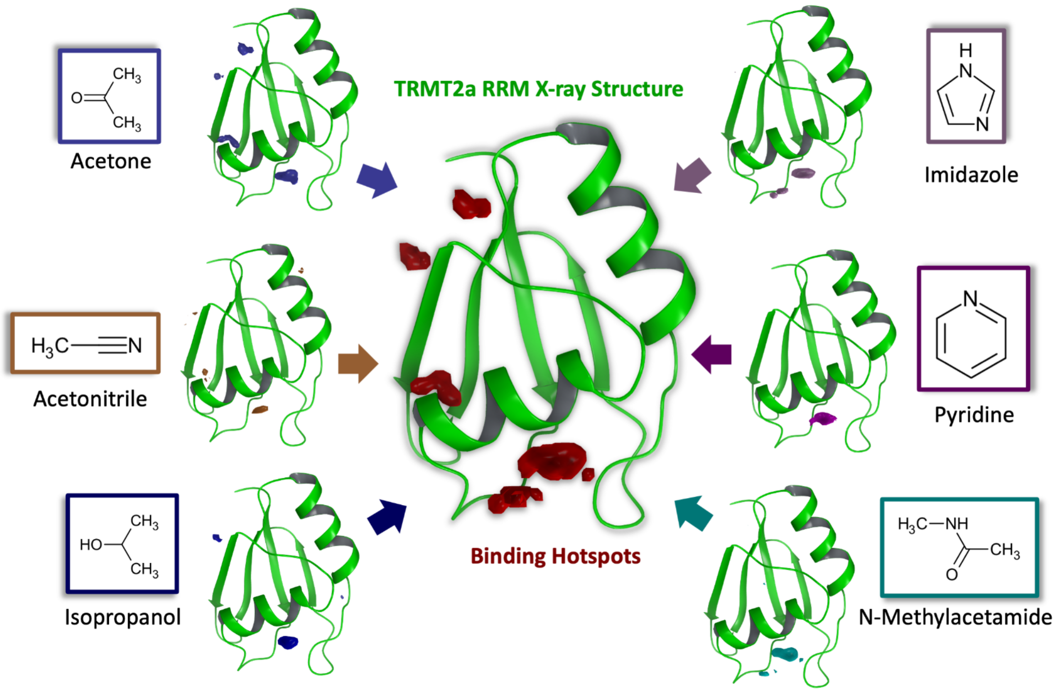

**Figure S16***.* Binding hotspot mapping with Mixed Solvent Molecular Dynamics of TRMT2A RRM. High propensity regions (contoured at an isovalue of 20σ) for individual cosolvent probes are color coded. All mixtures contain water and one type of small organic probes (binary solvent mixtures).

During these simulations, polar probes (acetone, acetonitrile, and isopropanol) localize on the surface of the β-sheet and contribute to the two hotspots (red) on the left top of the RRM (rear view orientation) depicted in the center. In contrast, all probes contribute to the hotspots on the helical back, while aromatic probes (imidazole and pyridine) exclusively contribute to them. The protein backbone-like cosolvent (N-methylacetamide) is also found preferentially in this area.

**TABLES**

**Table 1.** Statistical information of the X-ray experiments

| **Data collection** | | |
|---|---|---|
| PDB ID | 7NTN | 7NTO |
| Beamline | DESY P11 | SLS X06DA PXIII |
| Wavelength | 1.009300 | 1.000033 |
| Space group | *F*432 | *C*2 |
| Cell dimensions (Å, °) | a = b = c = 131.45 Å | a = 52.88, b = 36.30, c = 39.10, β = 119.18 |
| No. of molecules per asymmetric unit | 1 | 1 |
| Resolution (Å) | 50.00-2.02 (2.02-2.07) | 50.00–1.23 (1.23-1.26) |
| $R_{merge}$ | 5.800 (110.1) | 4.4 (105.8) |
| $I / \sigma I$ | 36.42 (3.85) | 18.2 (1.95) |
| CC (1/2) | 100.0 (92.0) | 100.0 (61.4) |
| Completeness (%) | 100.0 (100.0) | 97.0 (94.4 ) |
| Redundancy | 29.45 (31.92) | 6.62 (6.60) |
| **Refinement** | | |
| Resolution (Å) | 2.02 | 1.23 |
| No. reflections | 6,872 | 17,520 |
| $R_{work}$ / $R_{free}$ (%) | 20.51 / 25.14 | 12.71 / 17.15 |
| No. atoms | | |
| Protein | 648 | 708 |
| Water | 31 | 81 |
| Other atoms | 29 | 34 |
| *B*-factor overall | 41.66 | 18.83 |
| R.m.s. deviations | | |
| Bond lengths (Å) | 0.01 | 0.018 |
| Bond angles (°) | 0.98 | 2.13 |
| Ramachandran plot | | |
| Most favored (%) | 98.67 | 100 |
| Additional allowed (%) | 1.33 | 0 |

*Values in parentheses are for highest-resolution shell.

**Table S2**.
Selected compounds from the combined structure-based and ligand-based *in silico* approaches. IUPAC names, Canonical Smiles and PubChem CID (https://pubchem.ncbi.nlm.nih.gov) are provided. "*" indicate the compounds with detected toxicity in *in cell* experiments.

| **№** | **IUPAC Name (common name – when available)** | **Canonical smiles** | **PubChem (CID)** |
|---|---|---|---|
| 1 | 2-(piperidine-1-carbothioylsulfanyl)ethyl benzoate | O=C(OCCSC(=S)N1CCCCC1)C2=CC=CC=C2 | 343736 |

| | | | |
|---|---|---|---|
| 2 | (2Z,3S)-2-[(3,4,5-trimethoxyphenyl)methylidene]-1-azabicyclo[2.2.2]octan-3-ol | COC1=CC(=CC(=C1OC)OC)C=C2[CH](O)C3CCN2CC3 | 425101 |
| 3 | 4-chloro-N-[(4-methyl-5-...[2-(piperidin-1-yl)ethyl]sulfanyl}-4H-1,2,4-triazol-3-yl)methyl]aniline | CN1C(=NN=C1SCCN2CCCCC2)CNC3=CC=C(C=C3)Cl | 1156390 |
| 4 | (2S)-2-aminohexanoic acid (L-norleucine) | CCCCC(C(=O)O)N | 21236 |
| 5 | (4-carboxyphenyl)(chloro)mercury (PCMB) | C1=CC(=CC=C1C(=O)[O-])[Hg]Cl | 21226146 |
| 6 | 1-[(1R,2S,3aS,3bR,5aS,7S,8S,9aS,9bS,11aS)-1,7-bis(acetyloxy)-9a,11a-dimethyl-2-(1-methylpiperidin-1-ium-1-yl)-hexadecahydro-1H-cyclopenta[a]phenanthren-8-yl]-1-methylpiperidin-1-ium | CC(=O)OC1CC2CCC3C(C2(CC1[N+]4(CCCCC4)C)C)CCC5(C3CC(C5OC(=O)C)[N+]6(CCCCC6)C)C | 441289 |
| 7 | (2R,3R,4S,5S)-2-(6-amino-9H-purin-9-yl)-5-[(methylsulfanyl)methyl]oxolane-3,4-diol | CSCC1C(C(C(O1)N2C=NC3=C(N=CN=C32)N)O)O | 439176 |
| 8 | (2S)-2-amino-4-(… (2S,3S,4R,5R)-5-(6-amino-9H-purin-9-yl)-3,4-dihydroxyoxolan-2-yl]methyl}sulfanyl)butanoic acid | C1=NC(=C2C(=N1)N(C=N2)C3C(C(C(O3)CSCCC(C(=O)O)N)O)O)N | 439155 |
| 9 | (2R)-2-amino-4-(methylsulfanyl)butanoic acid (L-Methionine) | CSCCC(C(=O)O)N | 6137 |
| 10 | 2-amino-4-(ethylsulfanyl)butanoic acid (L-ethionine) | CCSCCC(C(=O)O)N | 6205 |
| 11 | butane-1,4-diamine (putrescine) | C(CCN)CN | 1045 |
| 12 | (4-aminobutyl)(3-aminopropyl)amine (spermidine) | C(CCNCCCN)CN | 1102 |
| 13 | 1-phenyl-4-(pyridine-3-carbonyl)piperazine | C1CN(CCN1C2=CC=CC=C2)C(=O)C3=CN=CC=C3 | 258236 |
| 14 | (2R)-2-[(2E)-3-phenylprop-2-en-1-yl]butanedioate | C1=CC=C(C=C1)C=CCC(CC(=O)O)C(=O)O | 348628 |
| 15 | ((2R)-4-acetyl-2-[(benzylsulfanyl)methyl]-5-oxo-1,2-dihydropyrrol-3-olate) | CC(=O)C1=C(C(NC1=O)CSCC2=CC=CC=C2)O | 54704354 |
| 16* | [4,6-bis(methoxycarbonyl)-5-(pyridine-2-carbonyl)pyrimidin-2-yl]sulfanide | COC(=O)C1=C(C(=NC(=S)N1)C(=O)OC)C(=O)C2=CC=CC=N2 | 3515120 |
| 17 | 2-{4-[5-bromo-2-(carboxymethoxy)benzoyl]phenoxy}acetate | C1=CC(=CC=C1C(=O)C2=C(C=CC(=C2)Br)OCC(=O)O)OCC(=O)O | 266989 |
| 18 | benzyl (1R,2R,6S,7S)-3,5-dioxo-4-azatricyclo[5.2.1.0^{2,6}]dec-8-en-4-yl carbonate | C1C2C=CC1C3C2C(=O)N(C3=O)OC(=O)OCC4=CC=CC=C4 | 93913 |
| 19* | 2-(phenylmethoxycarbonylamino)ethyl N-naphthalen-1-ylcarbamate | C1=CC=C(C=C1)COC(=O)NCCOC(=O)NC2=CC=CC3=CC=CC=C32 | 238681 |

| | | | |
|---|---|---|---|
| 20 | {4-oxo-2H,4H,5H-pyrazolo[3,4-d]pyrimidin-2-yl}methyl 2,2-dimethylpropanoate | CC(C)(C)C(=O)OCN1C=C2C(=O)NC=NC2=N1 | 135493751 |
| 21 | ethyl N-(benzyloxy)carbamate | CCOC(=O)NOCC1=CC=CC=C1 | 255288 |
| 22* | N-[(3-chlorophenyl)methyl]-4-hydroxy-2-(pyridin-2-yl)pyrimidine-5-carboxamide | C1=CC=NC(=C1)C2=NC=C(C(=O)N2)C(=O)NCC3=CC(=CC=C3)Cl | 70768840 |
| 23* | N'-benzyl-2-(4-chlorophenoxy)ethanimidamide | C1=CC=C(C=C1)CN=C(COC2=CC=C(C=C2)Cl)N | 4672872 |
| 24 | 1-(4-methoxyphenyl)-4-(pyridine-3-carbonyl)piperazine | COC1=CC=C(C=C1)N2CCN(CC2)C(=O)C3=CN=CC=C3 | 976616 |
| 25 | 3-(4-phenylpiperazine-1-carbonyl)pyridin-1-ium-1-olate | C1CN(CCN1C2=CC=CC=C2)C(=O)C3=C[N+](=CC=C3)[O-] | 17558279 |

### SUPPLEMENTAL TEXT

#### Supplements S1: additional REST simulations

Two replica exchanges with solute tempering (REST2) simulations[10, 11] were performed for 250 ns. The first one was started from a selected open conformation with respect to the distance of F92-W134 (discussed in the main text), while the second one started from the crystalized structure with the close conformation. Both simulations started from the 2 Å resolution crystal structure of the TRMT2A RRM from which the original MD was performed.
Classical MD simulations at different temperatures were performed for 50 ns spanning between 300 K to 520 K, in order to estimate the limit temperature where unfolding events happen (data not shown). For the open conformation, the limit was found to be around 400K, where the loops started to move drastically and the tertiary structure started to partially demise. Thus, the higher limit of 400 K was set for the REST2 simulations.

As the REST2 started from an open conformation, it is possible to explore different conformations with various degrees of cryptic pocket opening (Figure S9). However, the majority of the conformations are consistent with a partially close state, with the π-systems of F92 and W134 are either in a near parallel-displaced or perpendicular-like (T-shaped). The last 90 ns of REST2 simulations were used to build a conformational landscape using the ring-to-ring angle and distance (Figure S10). The projected conformational landscape suggests that those two parallel conformations are energetically similar, with a very small difference in favor of the flipped F92. The T-shaped conformation, which corresponds to the transition energy barrier between the two parallel conformations is estimated in the order of 2 $k_BT$.

On the other hand, in the REST2 starting from the closed conformation, an opening event is never found, consistent with the work of Oleinikovas *et. al*. [12], which indicates that the opening of the cryptic sites cannot be enhanced by pure temperature-based enhanced sampling. A difference of ~6.5 $k_BT$ between the closed conformation and the opening of the cryptic pocket was indeed estimated. Our findings suggest that the opening of the cryptic pocket is a rare event that might be facilitated by ligand binding but not by temperature or Hamiltonian scaling.

**Supplements S2**

Force field: OPLS3e; Ionization: Generate possible states at pH 7.0 +/- 2.0 using Epik; Desalt: yes; Generate tautomers: yes; Stereoisomers: Retain specific chiralities, Generate at most 32 per ligand; Output format: Maestro.

**Supplements S3**

Both compounds were defined as active; "Find best alignment and common features" was selected as pharmacophore method; Hypothesis Settings -> Features: Hypothesis should match at least 100% of actives; Number of features in the hypothesis: 4 to 5; Hypothesis difference criterion: 0.5; Minimum values of 0, maximum values of 3 and tolerance values of 2 were assigned to all features; 10 hypotheses should be generated. Hypothesis Settings -> Scoring: "Phase Hypo Score" was selected as scoring function; No excluded volume shell was created since the binding pose of compounds 13 and 15 could not be validated experimentally.

**Supplements S4**

The descriptors used to assess ADMET properties were the Predicted brain/blood partition coefficient, Predicted apparent MDCK cell permeability in nm/sec, Predicted aqueous solubility (log S), Number of reactive functional groups. In order to get an idea of the range in which the values of these descriptors should be for compounds that pass the filter, all structures of approved CNS drugs (beside anesthetics) listed in the DrugBank database (https://www.drugbank.com) were downloaded, and their descriptor values were calculated.
